# MT4-MMP-mediated NRP1 shedding fine-tunes VEGFA signaling dynamics during embryonic brain angiogenesis

**DOI:** 10.64898/2026.03.26.714199

**Authors:** Emma Muñoz-Sáez, Natalia Moracho, Cristina Clemente, Diana Cordón-Romero, Alberto Jiménez-Montiel, María Losa-Fontangordo, Rodrigo Torrillas-de la Cal, Juan Francisco Aranda, Guido Serini, Esther Serrano-Saiz, Emilio Camafeita, Jesús Vázquez, Fernando Martínez, Alicia G. Arroyo, Cristina Sánchez-Camacho

**Affiliations:** Universidad Europea de Madrid (UEM), Faculty of Biomedical and Health Sciences, Department of Biosciences, Madrid, Spain; Department of Biomedicine, Center for Biological Research Margarita Salas (CIB-CSIC), Madrid, Spain; Program of Myocardial Homeostasis & Cardiac Injury, Centro Nacional de Investigaciones Cardiovasculares (CNIC), Madrid, Spain; Department of Genetics, Physiology and Microbiology, Faculty of Biological Sciences, Complutense University of Madrid (UCM), Madrid, Spain; Centro de Biología Molecular Severo Ochoa (CBM, CSIC-UAM), Madrid, Spain; Escuela de Doctorado, Universidad Autónoma de Madrid, Madrid, Spain; Candiolo Cancer Institute, Fondazione del Piemonte per l’Oncologia (FPO) - IRCCS, Candiolo (TO), Italy; Department of Oncology, University of Torino, Candiolo (TO), Italy; Laboratory of Cardiovascular Proteomics, Centro Nacional de Investigaciones Cardiovasculares (CNIC), Madrid, Spain; CIBER de Enfermedades Cardiovasculares (CIBERCV), Madrid, Spain; Bioinformatics Unit, Centro Nacional de Investigaciones Cardiovasculares (CNIC), Madrid, Spain

**Keywords:** Neurovascular development, angiogenesis, MT4-MMP (MMP17), Neuropilin-1 (NRP1), VEGFA signaling, wound healing, vascular remodeling

## Abstract

Angiogenesis is essential for embryonic brain development and tissue repair, yet the mechanisms that spatiotemporally coordinate endothelial behavior to ensure balanced vascular remodeling remain elusive. Here, we identify the glycosylphosphatidylinositol (GPI)-anchored protease MT4-MMP as a critical, context-dependent determinant of angiogenic growth. Global loss of MT4-MMP transiently impairs vascular network formation in the embryonic hindbrain, whereas endothelial-specific deletion triggers an aberrant angiogenesis characterized by increased vessel density, branching, and a profound loss of vascular organization. This dual phenotype reveals MT4-MMP as a fundamental coordinator of neurovascular development. Consistently, MT4-MMP expression was dynamically regulated during wound repair, and its absence amplifies angiogenesis and accelerates wound closure in adult skin, highlighting its role in maintaining vascular homeostasis postnatally. Mechanistically, MT4-MMP-deficient endothelial cells exhibit impaired polarization and sustained, rather than transient, VEGFA-induced ERK activation. We identify NRP1 as a novel substrate of MT4-MMP and demonstrate that MT4-MMP-mediated NRP1 cleavage restricts NRP1 surface availability to tune the intensity of VEGFA signaling. Furthermore, pharmacological blockade of VEGFA–NRP1 binding partially rescues the vascular defects caused by endothelial MT4-MMP loss *in vivo*. Together, these findings uncover the MT4-MMP/NRP1 axis as a pivotal control point that prevents aberrant vessel expansion, establishing membrane-anchored proteolysis as a primary regulator across developmental and reparative contexts.

**Teaser:** The proteolytic constraint exerted by MT4-MMP dictates neurovascular development and wound repair through the spatial control of NRP1-VEGFA signaling.

## Introduction

Angiogenesis, the formation of new vessels from pre-existing ones, primarily occurs through capillary sprouting, leading to the expansion of the vascular networks during embryonic development. In the central nervous system (CNS), developmental vascularization involves a close crosstalk between endothelial cells and neural cells, which is not only critical for oxygenation but also relevant for subsequent brain-blood barrier (BBB) formation and prevention of brain pathologies ^1-3^. By E8.5 in the mouse, angioblasts are recruited to form the perineural vascular plexus (PNVP) around the neural tube, from which vessels sprout, ingress, and extend into the neural parenchyma, branching and fusing with neighboring radial vessels to form the subventricular vascular plexus (SVP) by E10.5-E12.5 ^4-6^. This intricate angiogenic process requires exquisite spatiotemporal orchestration to give rise to a stereotyped, mature vascular plexus capable to nurture the CNS ^5^.

During CNS vascularization, neural-derived VEGFA is absolutely required for vessel ingression from the PNVP ^7^. VEGFA includes soluble (VEGFA120) and heparin-bound (VEGFA165, VEGFA189) isoforms, whose expression and gradients must be precisely balanced. Thus, overexpression of VEGFA120 results in defective branching and vessel dilation, whereas that of VEGFA165/VEGFA189 leads to ectopic vessel entry points, excessive branching and thinner vessels in the nervous system of quail and mouse embryos ^6,7^. These matrix-associated VEGFA isoforms, particularly VEGFA165, can interact with the co-receptor neuropilin-1 (NRP1), a multidomain protein also able to bind class 3 semaphorins among other ligands ^8^, and essential for proper brain angiogenesis. Indeed, NRP1 mutants lacking expression globally or selectively in endothelial cells (but not in neural cells or macrophages), exhibit defective hindbrain vascularization ^9^. While NRP1 is clearly indispensable for later phases of subventricular plexus formation (SVP) formation, its precise regulation remains a subject of intense study. More precisely, NRP1 mutants show reduced and mis-patterned lateral branching, increased vessel size and failed anastomosis but not defects in vessel ingression into the neural tube ^10,11.^ Moreover, NRP1 actions in neural tube vascularization appeared to rely mainly on its regulation of VEGFR2 levels and activity rather than on VEGF binding ^12-14^, while NRP1 cytosolic tail signaling is relevant for arteriogenesis *in vivo* ^15^. NRP1 also participates in endothelial cell adhesion, integrin crosstalk and filopodia formation ^16-18^. Furthermore, unlike other cell types, NRP1 acts as a negative regulator of TGFβ specifically in endothelial cells *in vitro* ^19^. *In vivo,* endothelial NRP1 interacts with αVβ8 integrin expressed in the neuroepithelium or radial glia cells, thereby modulating TGFβ availability and signaling during CNS vascularization ^20^. Despite this extensive knowledge, it remains unclear how NRP1 actions are spatiotemporally fine-tuned to shape the brain vasculature during embryonic development.

There is growing interest in elucidating the regulation of embryonic brain angiogenesis due to its potential therapeutic implications in neurodegenerative diseases, which often involve vascular dysfunction. In this line, matrix metalloproteinases (MMP) are key regulators of vascular patterning, not only by degrading the extracellular matrix (ECM) but also by the release of angiogenic growth factors, the shedding of transmembrane proteins, and the activation/inhibition of soluble cytokines, etc. ^21^. Among these, MT4-MMP (MMP17), stands out as a glycosylphosphatidylinositol (GPI)-anchored protease expressed in both neural progenitors and scattered endothelial cells during CNS development, making it a good candidate to modulate neurovascular development ^22^. We had previously recognized MT4-MMP contribution to embryonic aorta development by promoting the differentiation of smooth muscle cells via osteopontin cleavage and JNK signaling ^23^. However, whether this membrane-anchored protease plays any role in endothelial cells and during developmental angiogenesis remains entirely unknown.

In this study, we identify MT4-MMP as a crucial, context-dependent modulator of embryonic brain vascularization. Using endothelial cell-specific conditional mice, we demonstrate that MT4-MMP is essential for fine-tuned and balanced angiogenesis, as its absence triggers an enhanced and disorganized angiogenic response. This regulatory role is recapitulated during adult wound-healing, where MT4-MMP deficiency results in exacerbated angiogenesis and accelerated wound closure. Mechanistically, we uncover a novel proteolytic axis in which MT4-MMP cleaves NRP1, thereby limiting its surface availability and tuning the VEGFA-ERK signaling, essential for proper CNS vascularization. Our findings establish the MT4-MMP/NRP1 axis as a fundamental control point for vascular homeostasis with broad implications for CNS development, tissue repair and vascular diseases.

## Materials and Methods

### Mice

C57BL/6 wild-type mice were purchased from Charles River Laboratories (Spain). The *Mt4-mmp-*LacZ (*Mt4-mmp*^LacZ/+^) reporter mice and the homozygous *Mt4-mmp*^LacZ/LacZ^ (*Mt4-mmp^-/-^*) mice maintained in the C57BL/6 background were previously described ^24^. Pregnant females were sacrificed by cervical dislocation, and embryonic hindbrains were dissected at embryonic days (E)10.5, E11.5 and E12.5 (with E0.5 defined as the day of vaginal plug detection). *Mt4-mmp*^loxP/loxP^ mice were generated in our lab and, after backcrossing into the C57BL/6 background, were crossed with *Cdh5-*Cre^ERT2^ mice (kindly provided by Prof. R. Adams’ laboratory) to generate *Mt4-mmp*^loxP/loxP^; *Cdh5-*Cre^ERT2/+^ mice (*Mt4-mmp^f/f^* and *Mt4-mmp^iΔEC^* mice). Mice were housed and all animal experiments were performed under specific pathogen-free conditions at animal facilities of the Centro de Investigaciones Biológicas Margarita Salas (CIB-CSIC), and in strict accordance with the institutional guidelines. Mice were kept under a 12-h light/dark cycle (lights on from 07:00 to 19:00 h). *Mt4-mmp*^LacZ/+^ embryos were genotyped by tail DNA extraction and PCR using the kit REDExtract-N-Amp™ Tissue PCR Kit (254-457-8, Sigma) and the following primers were used for LacZ, 5’-TCAGACACAGCCAGATCAGG-3’, 5’-AGCAACACGGCATCCACTAC-3’ and 5’-AATATGCGAAGTGGACCTGG-3’. Embryos from *Mt4-mmp*^loxP/loxP^; *Cdh5-*Cre^ERT2/+^ mouse crosses were genotyped by heart DNA extraction and PCR using the following primers for *Mt4-mmp*, 5’-CCTAATGTACATAGCCAGCCAAG-3’ and 5’-AAGGCCAGGTGTGTTCAATC-3’; and for Cre recombinase, 5’-AGGTGTAGAGAAGGCACTTAGC-3’ and 5’-CTAATCGCCATCTTCCAGCAGG-3.

### Administration of tamoxifen and NRP1 inhibitor

Tamoxifen (T5648, Merck) was dissolved at 30 mg/ml in 10% ethanol/90% corn oil. Pregnant females received intraperitoneal (i.p.) injections of tamoxifen (2 mg/30 g body weight) daily for three consecutive days corresponding to E7.5, E8.5 and E9.5 In parallel experiments, the NRP1 inhibitor EG00229 trifluoroacetate (MedChem Express, HY-20799) was dissolved at 20 mg/ml in DMSO and administered i.p. to pregnant females at a dose of 10 mg/kg at E9.5 and E10.5; control pregnant females were injected with the corresponding vehicle alone. Embryonic hindbrains were dissected at E11.5 and E12.5 for immunofluorescence analysis.

### LacZ staining

*Mt4-mmp*^LacZ^ embryos isolated at E11.5 were fixed in 0.125% glutaraldehyde for 40 minutes at 4 °C. Subsequently, LacZ staining was performed according to the protocol described in Blanco *et al*., 2018 ^25^. Paraffin-embedded *Mt4-mmp*^LacZ^ embryos were sectioned at 8 µm.

### Whole-mount and section immunofluorescence

Whole-mount hindbrain staining was performed as described by Fantin *et al.*, 2013 ^26^. Hindbrains were incubated with biotinylated isolectin B4 (IB4, L2140, Sigma; 1:200) and anti-ERG-Alexa647® (ab196149, Abcam; 1:500) in blocking buffer (10% NGS in PBS with 0,2% Triton X-100) for 48 hours at 4 °C, followed by streptavidin-Alexa594® (1:100) incubation overnight at 4°C. Flat-mount hindbrains were processed as described^26^. Representative images were acquired on a Leica SP8 confocal microscope with a 20x objective and z-stacks were captured every 1.5 µm. For section immunohistochemistry, 10-15 µm cryosections were obtained from 4% paraformaldehyde (PFA)-fixed hindbrains. After blocking with mouse Fc Block^TM^ 2.4G2 antibody (BD Biosciences, 553142), sections were stained for IB4 (1:500), MT4-MMP (LEM-3/10 mouse IgG1 monoclonal antibody generated in our laboratory, 25 μg/ml), mouse isotype IgG1 control (eBioscience, 16-4714-85, 25 μg/ml), NRP1 (A13G13, Selleckchem; 1:100 or AF566, R & D; 10 μg/ml), pERK (9101, Cell Signaling, 1:100) and Hoechst 33342 (H1399, Invitrogen; 1:1000). Images were acquired with a 40x objective (z-stacks interval: 1 µm).

### Image quantification

For each hindbrain, maximum intensity projections (MIP) from four independent fields (two per each side of the floor plate) were analyzed. Vascular parameters were quantified in two independent 0.25 mm^2^ regions of interest (ROIs) per field using AngioTool v0.6q and following the protocol previously described ^27^. Vascular pERK abundance was quantified in neural hindbrain vessels using Fiji/ImageJ ^28^. Briefly, ROIs were defined around the neural vasculature, excluding the perineural vascular plexus (PNVP). Vasculature was selected based on IB4 signal using a default threshold and automatic removal of particles < 1 µm^2^. The pERK-positive area was segmented after applying a default threshold for wild-type and MT4-MMP-null hindbrains or a Gaussian filter (sigma=0.5) and an Otsu threshold for embryonic hindbrains from *Mt4-mmp*^f/f^ and *Mt4-mmp*^iΔEC^ mice. Results are expressed as the percentage of pERK+ area within the IB4 mask.

### *In situ* hybridization

E11.5 *Cx3cr1*^GFP/+^ ^29^ embryos were fixed for 2 h at room temperature (RT) in 4% PFA and cryoprotected in 15% sucrose in PBS. Embryos were embedded in 7.5% gelatin (G2625, Sigma)/15% sucrose and frozen at −80 °C. Horizontal cryostat sections (16 μm thickness) were collected and hybridized with digoxigenin-labeled riboprobes as described ^30^. A mouse *Mt4-mmp* 670 bp riboprobe was generated using the following primers: Fw: 5’-TGTACTGGCGCTATGATGACCAC−3’ and Rv: 5’-GTGCAACGAAGCAGCATGATGT−3’. The rat *Nrp1* riboprobe was previously described ^31^. Sections were then stained for ERG, and images were acquired with a 40x objective (z-step: 1 µm).

### Binding assays with AP-VEGF ligand

VEGFA165-AP binding assays were performed as previously described ^26^. Briefly, E11.5 hindbrains were dissected on ice-cold PBS, and the meninges/mesenchyme were removed to expose the ventricular surface. Tissue was blocked in 10% fetal bovine serum (FBS) in PBS for 30 minutes at 4 °C. Flat-mount hindbrains were incubated with VEGFA_165_-AP in conditioned medium overnight at 4 °C and then fixed in 4% PFA at 4 °C for 30 min. Endogenous alkaline phosphatase activity was inactivated by incubation in PBS for 30 minutes at 65 °C. Finally, samples were incubated in AP-development buffer (100 mM Tris-HCl pH 9.5, 100 mM NaCl, 5 mM MgCl_2_) with NBT/BCIP substrate until signal developed (∼20-30 min). AP-only controls assessed background. Samples were imaged by bright-field microscopy.

### Wound healing assay

The backs of wild type, *Mt4-mmp^-/-^, Mt4-mmp^f/f^* and *Mt4-mmp^iΔEC^* mice were shaved at day -1. At day 0, four full-thickness wounds were performed with a punch in the back area. Pictures were taken for 9 days with a ruler scale for normalization. The kinetics of wound closure was quantified as the remaining wound area (% of initial area) using Fiji/ImageJ.

### Proteomics analysis

Skin tissue was harvest from around the wound site 6 days post-lesion from wild-type and MT4-MMP-null mice, immediately frozen in liquid nitrogen, and stored at −80°C. A total of 10 samples was analyzed (comprising 5 pools of mice per genotype). Protein extracts were obtained by mechanical homogenization in FastPrep tubes using a lysis buffer (50 mM Tris-HCl pH7.5, and 2% SDS) with three rupture cycles (6,000 rpm for 1 min each), followed by centrifugation at 12,000 x g for 10 min. The resulting protein extracts were quantified using the RC DC protein assay (Bio-Rad). Proteins (100 μg) were on-filter digested using FASP tubes (Expedeon), and the resulting peptides were isobarically labelled using TMT10plex reagents (Thermo Scientific) according to the manufacturer’s instructions. The labelled peptides were mixed, and the resulting sample was desalted using HLB cartridges (Oasis, Waters Corporation, Milford, MA, USA). To increase proteome coverage, peptides were fractionated into six fractions on C18 reversed-phase columns (High pH Fractionation Kit; Thermo Scientific) following the supplier’s protocol. The resulting labelled peptide samples were subjected to liquid chromatography-tandem mass spectrometry (LC-MS/MS) analysis on an Easy nLC 1000 nano-HPLC system (Thermo Scientific) coupled to a Q Exactive Orbitrap mass spectrometer (Thermo Scientific). Peptides were resuspended in Buffer A (0.1% formic acid) and loaded onto a pre-column (PepMap100 C18 LC, 75 μm ID, 2 cm; Thermo Scientific). On-line separation was performed on a NanoViper PepMap100 C18 LC analytical column (75 μm ID, 50 cm; Thermo Scientific) using a continuous gradient from 5%–32% B for 240 min and 32%–90% B for 5 min (B = 100% acetonitrile, 0.1% formic acid) at a flow rate of 200 nL/min. Each MS run consisted of full-scan FT-resolution spectra (60,000FWHM) in the 400–1,500 *m/z* range, followed by data-dependent MS/MS spectra of the most intense parent ions. The AGC target value for the Orbitrap was set to 200,000. Fragmentation was performed at 33% normalized collision energy (NCE) with a target value of 50,000 ions, 30,000 resolution, 120 ms injection time, and 45 s dynamic exclusion. For peptide identification, MS/MS spectra were searched using the Sequest HT algorithm implemented in Proteome Discoverer (v1.4; Thermo Fisher Scientific) against the *Mus musculus* Uniprot database (January 2015 release; 16,715 entries). The search parameters were set as follows: trypsin digestion with up two missed cleavage sites; precursor and fragment mass tolerances of 2 Da and 0.02 Da, respectively; carbamidomethylation of cysteine and TMT modification at the N-terminous and lysine residues as fixed modifications and methionine oxidation as a dynamic modification. Peptide identification was performed using the probability ratio method ^32^. The false discovery rate (FDR) was calculated using an inverted (decoy) database and the refined method ^33^, applying a 1% FDR cut-off and an additional filtering step with a precursor mass tolerance of 15 ppm ^34^. Statistical assessment of differential protein abundance was conducted usingthe Weighted Spectrum, Peptide and Protein (WSPP) statistical model ^35^ and the SanXoT package ^36^. Functional category analysis of coordinated protein behaviour was performed based on the Systems Biology Triangle model ^37^. Functional categories were retrieved from Gene Ontology (GO), Ingenuity Pathway Analysis (IPA), KEGG, and DAVID databases.

### Skin Flow cytometry

Skin was collected 6 days post-injury using a punch in the wound and a larger punch in the periphery. Tissues were disaggregated by incubation in collagenase and blocked in PBS containing 5% BSA and anti-CD16/CD32 (2.4G2, BD Pharmingen, 553142; 1:100) for 15 min at 4 °C. Samples were stained for 30 min at 4 °C with the primary antibodies anti-CD45-V450 (48-0451-82, eBioscience) and anti-CD31-FITC (553372, BD Bioscience) and with anti-mouse NRP1 (AF566-SP, R&D) and a secondary anti-goat-Alexa568® antibody. Data were acquired on a FACSCanto III cytometer (BD) and analyzed using FlowJo software (BD Biosciences).

### Mouse Aortic Endothelial Cell (MAEC) culture

MAECs were isolated from aortas of 4-week-old mice. Aortas were incubated for 5 min at 37°C in 0.2% collagenase type I (LS004194, Worthington), to allow removal of the adventitia. Aortas were cut into small pieces (1–2 mm) and digested for 45 min at 37 °C in DMEM containing 6 mg/ml collagenase type I and 2.5 mg/ml elastase (LS002290, Worthington). EC colonies were purified by two rounds of positive selection with anti-ICAM2 and magnetic beads. MAECs were cultured on 0.2% gelatin-coated plates in DMEM/F12 supplemented with 20% FBS, 100 U/ml penicillin, 100 μg/ml streptomycin, 2 mM L-glutamine, 10 mM HEPES, and ECGS/heparin

### Endothelial cell polarization and migration in scratch assays

Confluent MAEC monolayers were serum-starved for 2 hours and scratched with a plastic pipette tip and washed with HBSS. Cells were cultured with or without 20 ng/ml of VEGFA. For collective migration, time-series pictures were recorded every 20 min overnight, and motility parameters were analyzed with the Ibidi Chemotaxis tool (ImageJ). For polarization, cells were fixed in 4% PFA for 15 min at RT, blocked, and stained with the primary antibodies rabbit anti-ERG-Alexa647® (Ab196149, Abcam, endothelial cell nucleus) and mouse anti-GM130 (610823, BD Biosciences, Golgi) overnight in a cold room, followed by the corresponding anti-mouse secondary antibody. Cell orientation was quantified as the nucleus–Golgi axis angle relative to a line perpendicular to the wound edge.

### Protein extraction and Western-blot analysis

MAECs were serum-starved and stimulated with 20 ng/ml VEGFA for different times (from 5 up to 60 minutes). After washing, cells were lysed in RIPA buffer supplemented with protease and phosphatase inhibitors (11697498001, Roche). Protein samples were separated by 10% SDS–PAGE, transferred to nitrocellulose membranes (1620115, Bio Rad), blocked with 5% BSA, and incubated with the primary antibodies: anti-pPLCβ3/PLCβ3 (2481 and 14247 Cell Signalling, 1:1000), anti-pERK/ERK (9101 and 9102 Cell Signaling, 1:1000), and anti-pp38MAPK/p38 MAPK (9211 Cell Signaling, 1:1000; MA5-15116 Invitrogen, 1:1000). After several washes, the membrane was incubated with the corresponding horseradish peroxidase (HRP)-conjugated secondary antibodies. Bound secondary antibodies were visualized with Luminata Classico Western HRP Substrate in ChemiDoc Imaging System (ThermoFisher Scientific). Western blots were quantified using ImageJ software.

For NRP1 shedding assays, supernatant and lysates from Bone Marrow-Derived Monocytes (BMDMo) from wild-type and MT4-MMP-deficient mice were collected before lysis in RIPA buffer supplemented with protease and phosphatase inhibitors (11697498001, Roche). Protein samples were separated by 10% SDS–PAGE, transferred to nitrocellulose membranes (1620115, Bio Rad), blocked with 5% BSA and incubated sequentially with the primary antibodies anti-NRP1 (AF566-SP, R&D) and anti-tubulin (T6074, Sigma-Aldrich). After several washes, the membrane was incubated with the corresponding horseradish peroxidase (HRP)-conjugated secondary antibodies. Bound secondary antibodies were visualized with Luminata Classico Western HRP Substrate in ChemiDoc Imaging System (ThermoFisher Scientific). Western blots were quantified using ImageJ software.

MAECs were serum-starved and stimulated with 20 ng/ml VEGF for 15 min before lysis in a buffer containing 10 mM Tris-HCl pH 7.5, 1% (w/v) Triton X-114, 150 mM NaCl, and protease inhibitors; samples were heated for 5 min at 30 °C to separate hydrophilic and lipophilic phases. Proteins were separated by 10% SDS-polyacrylamide gel electrophoresis, transferred to nitrocellulose membranes, and blocked with 5% non-fat milk. Primary antibodies used were anti-NRP1 (AF566-SP, R&D) and anti-tubulin (T6074, Sigma-Aldrich). After overnight incubation at 4 °C and washes, bound primary antibody was detected by incubation for 1 h with donkey-anti-rabbit (IRDyeTM 800CW or 680CW, Odyssey, 1:10,000) or donkey-anti-mouse (IRDyeTM 800CW or 680CW, Odyssey, 1:10,000) secondary antibodies, followed by visualization with the Odyssey Infrared Imaging System (LI-COR Biosciences).

### Co-transfection of HEK293 cells

The murine full-length Neuropilin-1 (mNRP1) cDNA kindly provided by Andreas Püschel (University of Münster, Germany), was cloned in frame into a pEGFP-N1 mammalian expression plasmid to create a C-terminally EGFP-tagged mNRP1 (mNRP1-EGFP). HEK293 cells were co-transfected over 3 hours with wild-type or catalytically inactive E248A-mutated mouse MT4-MMP vectors and the mNRP1-EGFP plasmid (3:1 ratio) using Lipofectamine 2000 (Invitrogen). After 48 hours, HEK293 cells were stained for flow cytometry with anti-MT4-MMP (AB39708, Abcam) and a secondary anti-rabbit-Alexa568® and with anti-mNRP1 (AF566-SP, R&D) and a secondary anti-goat-Alexa647®. The data were obtained using a FACSCanto III flow cytometer (BD). The geometric mean fluorescence intensity and median fluorescence intensity of NRP1 expression on cell surface of co-transfected HEK293 cells that were positive for both MT4-MMP and NRP1-EGFP were analyzed using FlowJo.

### *In vitro* digestion assay

Human recombinant NRP1-IgG1 Fc chimera (5 pmol) (or 500 ng) (10455-N1/CF, R&D) was incubated with the catalytic domain of human recombinant MT4-MMP (hrMMP17, P4928, Abnova; 5 or 10 pmol (or 250 ng) in digestion buffer (50 mM Tris-HCl, 1-10 mM CaCl_2_, 80 mM NaCl, pH7.4) for 2-3 hours at 37 °C. Samples were separated by 12% SDS-PAGE and transferred to nitrocellulose membranes (Bio-Rad). Full-length rhNRP1-IgG1 Fc and C-terminal fragments were detected using goat anti-human IgG1 antibody or with sheep anti-human NRP1 (AF3870-SP, R&D) and a rabbit-anti-goat or a donkey-anti-sheep secondary antibody-HRP. Bands were visualized with Immobilon® Classico Western HRP substrate (WBLUC0500, Merck).

### *In silico* modeling and molecular dynamics

Structural modeling of the human NRP1/MT4-MMP complex was performed using I-Tasser v5.1 and Rosetta v3.8. Molecular dynamics (MD) simulations were executed in Gromacs 2024.2 using an asymmetric mammalian plasma membrane (PMM) environment. A total production of 500 ns was sampled across three replicas.

The FASTA sequence of the mature human NRP1 protein (Uniprot ID: O14786, residues 22-923) was submitted to a local implementation of the I-TASSER software suite (v5.1) ^38^ for threading modeling. The best model, defined by minimal energy and correct folding (optimal structural alignment to templates), was selected. As this model lacked a correct folding of the transmembrane and intracellular domains (TMIC), the angles and secondary structure of the TM were manually modified according to the PSIPRED predictions. The resulting model was used as a template for TM remodeling and refinement using the *Relax* tool ^39,40^ of the Rosetta suite (v3.8, www.rosettacommons.org). The model with the lowest energy and a topology compatible with a transmembrane protein was selected as the final NRP1 model. For membrane positioning, the final model was submitted to the OPM server (http://opm.phar.umich.edu).

For complex docking, the previously modeled MT4-MMP ^23^ and the NRP1 model were positioned close to each putative cleavage site using PyMOL (v1.8). This dimeric model served as an initial template to generate complex models using the MP_Dockapplication ^41^ from the Rosetta Membrane framework (v3.8) and subsequently clustered. The model with the minimal energy (E) and a topology compatible with the cleavage of NRP1 by MT4-MMP (active site of MT4-MMP in the proximity to the predicted NRP1 cleavage site) was selected as the final model.

### Molecular Dynamics (MD) simulations for *in silico* models of mouse MT4-NRP1 complex modelled

The system setup was performed using a local implementation of GROMACS ^42-44^ (V2024.2) in two sequential steps: PMM membrane composition and PMM membrane optimization with PS/PC exchange between leaflets. Briefly, for the normal condition, the initial system (comprising the model, membrane ions and water) was created using the CHARMM-GUIweb server ^45^ with the following parameters:

|  | **model + lig** |
| --- | --- |
| OXTYPE | RECT |
| XTLTYPE | TETRAGONAL |
| A | 186.243845 |
| B | 186.243845 |
| C | 177.751 |
| ALPHA | 90 |
| BETA | 90 |
| GAMMA | 90 |
| ZCEN | 30.2885 |
| NLIPTOP | 642 |
| NLIPBOT | 600 |
| NWATER | 99.164 |
| WBOXZ | 22.5 |
| ISLCONJ | YES |
| ISMEMB | 1 |
| NCHARGE | -151 |

In all cases, a KCl concentration of 0.15 M and a temperature of 310.15 °K were maintained. The asymmetric mammalian plasma membrane (PMM) composition was defined as follows:

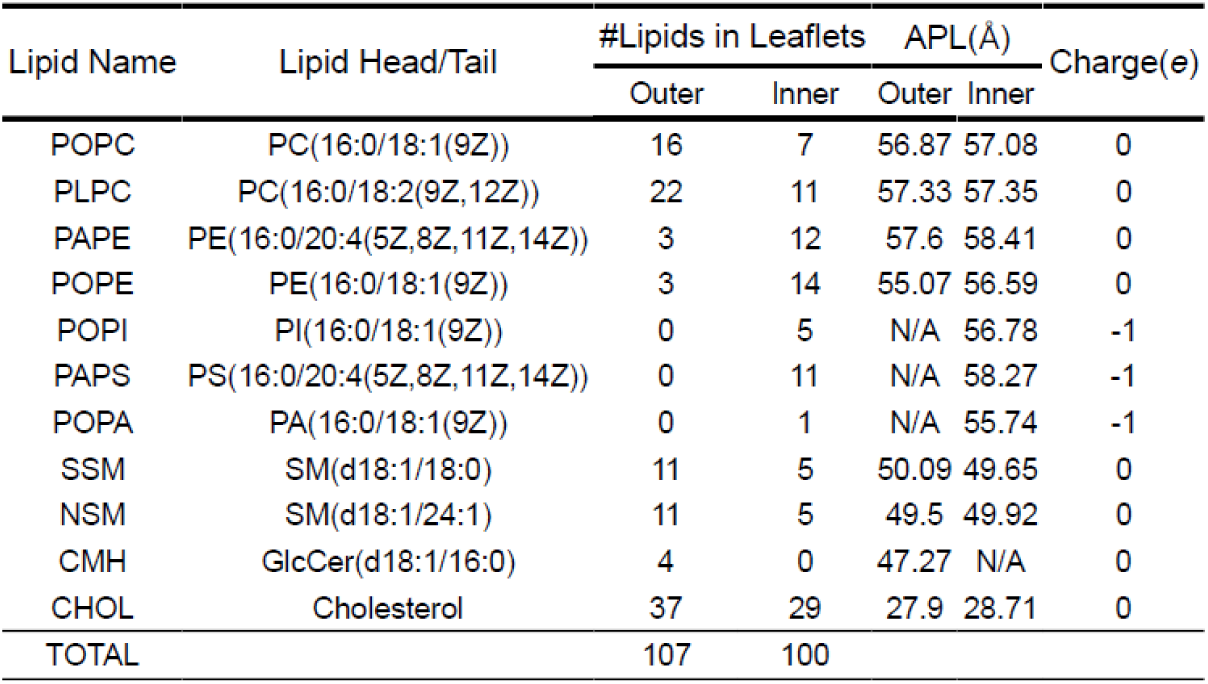

Starting from the initial input, each MD simulation was performed in GROMACS. An initial NPT minimization (constant number of particles, pressure at 1 atm, and temperature 310.15 °K) was conducted for 5,000 ps followed by six equilibration steps in the NPT ensemble (125,000 ps total). Subsequently, nine production steps of 500,000 steps each (with a time step *dt* = 0.002 ps) were executed and concatenated to obtain a final MD production trajectory of 500 ns (sampling interval 0.002 ps, compression ratio 1:10):

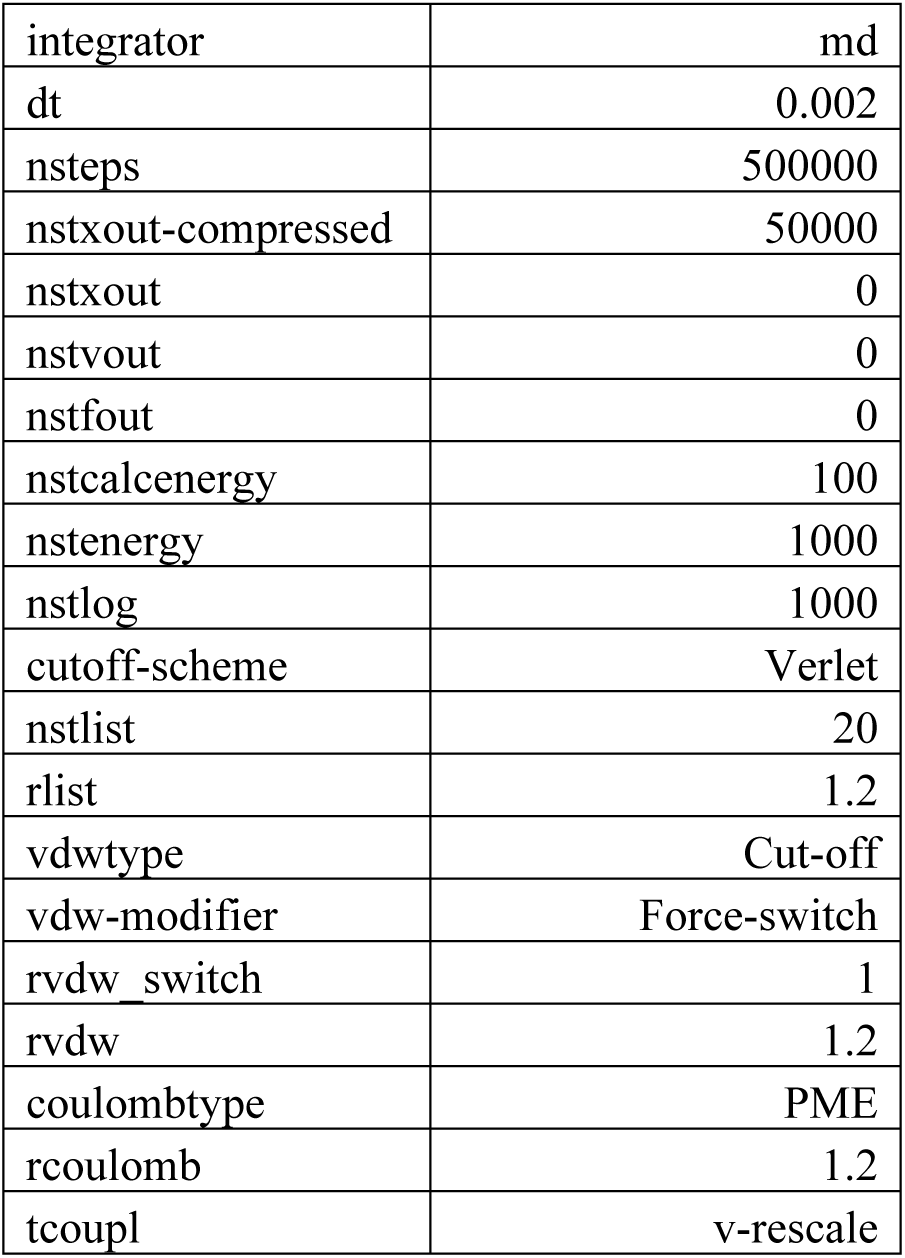
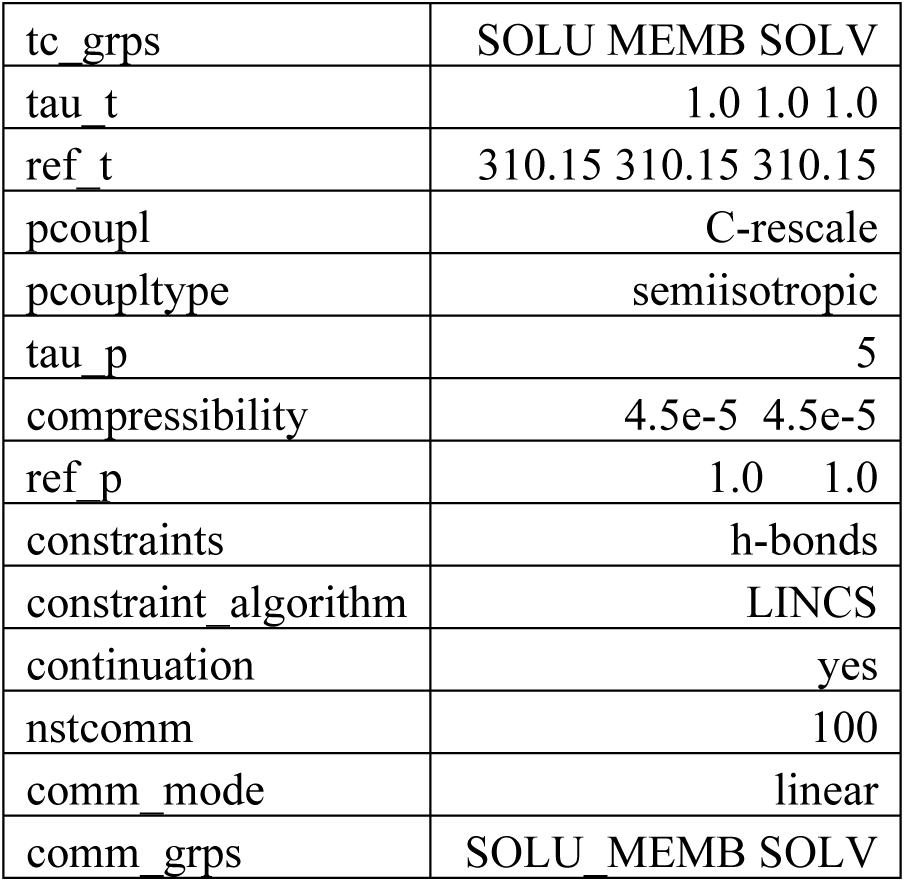

|  |  |
| --- | --- |
| integrator | md |
| dt | 0.002 |
| nsteps | 500000 |
| nstxout-compressed | 50000 |
| nstxout | 0 |
| nstvout | 0 |
| nstfout | 0 |
| nstcalcenergy | 100 |
| nstenergy | 1000 |
| nstlog | 1000 |
| cutoff-scheme | Verlet |
| nstlist | 20 |
| rlist | 1.2 |
| vdwtype | Cut-off |
| vdw-modifier | Force-switch |
| rvdw switch | 1 |
| rvdw | 1.2 |
| coulombtype | PME |
| rcoulomb | 1.2 |
| tcoupl | v-rescale |
| tc_grps | SOLU MEMB SOLV |
| tau_t | 1.0 1.0 1.0 |
| ref_t | 310.15 310.15 310.15 |
| pcoupl | C-rescale |
| pcoupltype | semiisotropic |
| tau_p | 5 |
| compressibility | 4.5e-5 4.5e-5 |
| ref_p | 1.0 1.0 |
| constraints | h-bonds |
| constraint_algorithm | LINCS |
| continuation | yes |
| nstcomm | 100 |
| comm_mode | linear |
| comm_grps | SOLU MEMB SOLV |

For each condition, three independent replicates were performed. The final state of the “Normal” condition served as the starting configuration for a new simulation with an altered membrane composition. This modification involved an exchange of phosphoadenosine 5’-phosphosulfate (PAPS) and phosphocholine (POPC) between leaflets, specifically incorporating PAPS into the extracellular (EC) leaflet.

|  |  |  |
| --- | --- | --- |
|  | EC | IC |
| POPC | 14 | 9 |
| PAPS | 2 | 9 |

The resulting lipid distribution was as follows:

|  |  |
| --- | --- |
|  | **model + lig** |
| BOXTYPE | RECT |
| XTLTYPE | TETRAGONAL |
| A | 206.882304 |
| B | 206.882304 |
| C | 183.417 |
| ALPHA | 90 |
| BETA | 90 |
| GAMMA | 90 |
| ZCEN | 25.8535 |
| NLIPTOP | 749 |
| NLIPBOT | 800 |
| NWATER | 97.562 |
| WBOXZ | 22.5 |
| ISLCONJ | YES |
| ISMEMB | 1 |
| NCHARGE | -189 |

MD simulations were performed as described above, with a final production of 500,000 steps per run (*dt* = 0.002 ps, compression ratio 1:10). For each condition, three independent replicates were conducted. Trajectory extraction and analysis were performed using the GROMACS command-line interface (CLI) and the MDAnalysis (v2.9.0) library.

### Statistics

Data collected were tested for normal distribution by Shapiro-Wilk test and compared with non-parametric or parametric statistical tests as appropriate (see figure legends for details). Graphs were prepared and statistical analysis performed using GraphPad Prism v11.0.0 (GraphPad Software). Differences were considered significant at: *p < 0.05; **p < 0.01; *** p < 0.001 and ****p < 0.0001.

### Study approval

Animal procedures were approved by the corresponding legal authority of the local government (Permit Number: PROEX 095.6/22). Animal studies were conformed to Directive 2010/63EU and recommendation 2007/526/EC regarding the protection of animals used for experimental and other scientific purposes, enforced in Spanish law under RD1201/2005.

## RESULTS

### MT4-MMP is transiently required for proper angiogenesis in the embryonic hindbrain

We investigated the role of MT4-MMP during hindbrain angiogenesis, based on our previous observations in *Mt4-mmp*^LacZ/+^ reporter mice ^22^ showing endothelial-specific transcription in the perineural vascular plexus (PNVP) at E9.5. Extending this analysis to later developmental stages, we detected *Mt4-mmp* (β-gal) expression not only in PNVP endothelial cells but also in vessels invading the neural parenchyma at E10.5 and E11.5 (**Supp. Fig. S1A-B**).

To determine the functional relevance of MT4-MMP, we assessed whether the absence of this protease may have any impact on developmental brain angiogenesis in *Mt4-mmp*^-/-^ embryos. Whole-mount staining of the endothelial marker IB4 at E11.5 revealed that the vascular network development was profoundly disrupted in the absence of MT4-MMP. Mutant hindbrains exhibited a hypovascular phenotype, characterized by significantly reduced vascular density and total vessel length along with narrower vessels in the subventricular vascular plexus (SVP) compared to wild type littermates (**Fig. 1A-B**). In addition, vascular connectivity was decreased, as evidenced by fewer junctions and increased lacunarity (space not occupied by blood vessels) in the SVP, indicating impaired network complexity (**Fig. 1B**). Notably, SVP vascular defects in *Mt4-mmp*^-/-^embryos were developmentally restricted to E11.5, revealing a transient, stage-specific requirement for MT4-MMP during the initial phase of neural parenchyma invasion (**Fig. 1C**).

**Figure 1.**
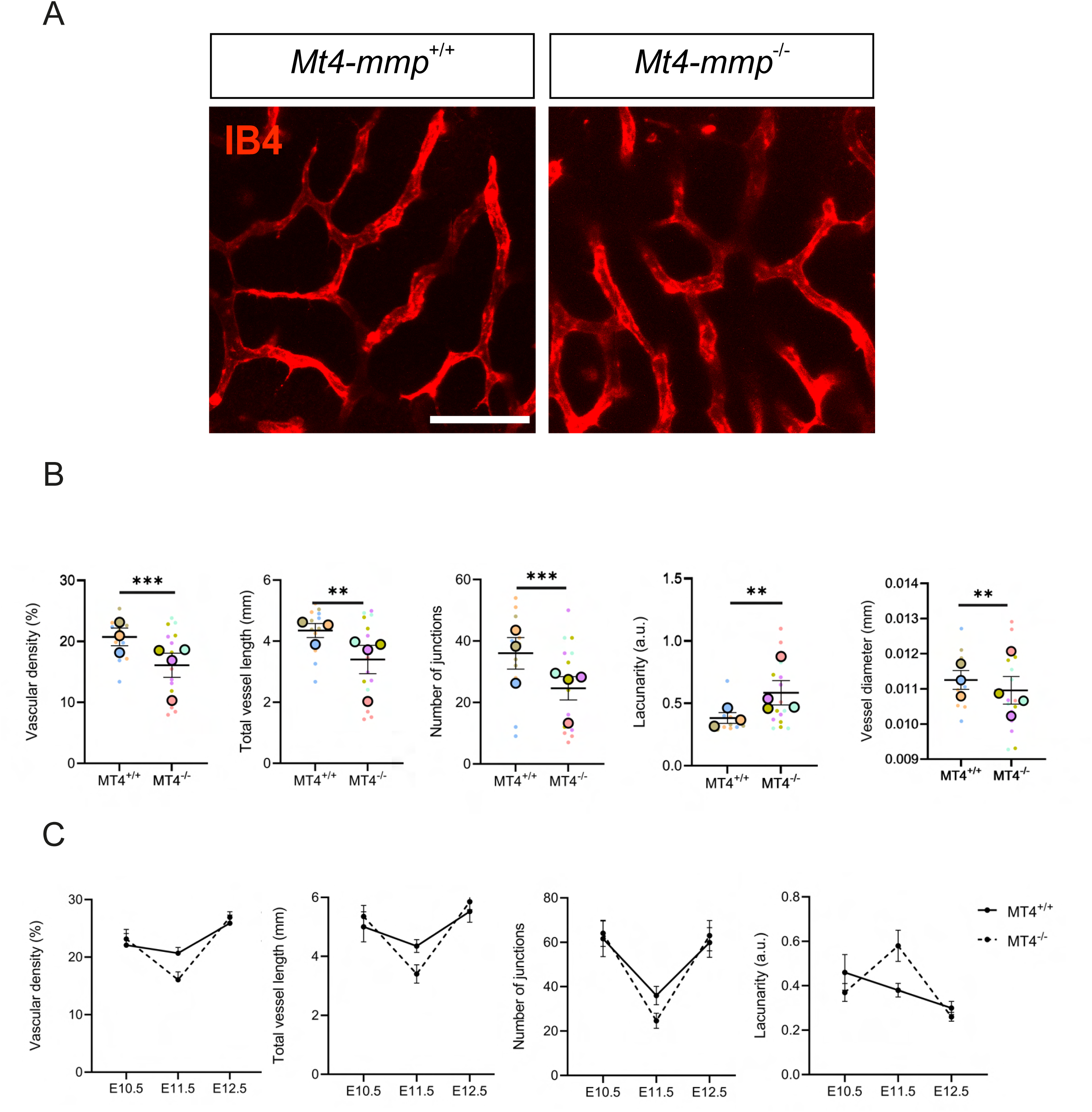
Angiogenesis is transiently impaired in the developing hindbrain of *Mt4-mmp*-null embryos. **A)** Representative maximum intensity projection (MIP) images of **whole-mount** E11.5 hindbrains from *Mt4-mmp*^+/+^ and *Mt4-mmp*^-/-^ embryos stained with IB4 (red); view shows the subventricular plexus (SVP). **B)** Quantification of vascular parameters (vascular density, total vessel length, number of junctions, and lacunarity) with AngioTool in the SVP of E11.5 hindbrains from *Mt4-mmp*^+/+^ (n=3) and *Mt4-mmp*^-/-^ (n=4) embryos. All the vascular parameters measurements refer to two different 0.25 mm^2^ crops per four hindbrain fields acquired in the confocal microscope. Average vessel diameter was estimated as the ratio of vascular area to vessel length. Values per individual image and the means ± SEM per embryo are shown in different colors. Data were analyzed using unpaired Welch’s t-test for parametric samples. **C)** Vascular parameters of the SVP in E10.5 to E12.5 hindbrains from *Mt4-mmp*^+/+^ and *Mt4-mmp*^-/-^ mice analyzed with AngioTool; means ± SEM are shown (n=3 embryos per stage and genotype). p-value <0.01 **, <0.001 ***. Scale bar:100 µm.

### Endothelial MT4-MMP fine-tunes vascular growth in a stage-dependent manner

MT4-MMP is expressed not only in endothelial cells but in other cell types, such as neural progenitors, that may influence angiogenesis during brain development ^22^. To dissect the cell-specific contribution of the protease, we generated inducible endothelial knockout embryos (*Mt4-mmp^i^*^ΔEC^) by administering tamoxifen via intraperitoneal injections to pregnant females (from E7.5 to E9.5) from *Mt4-mmp^f/f^*; *Cdh5-CreERT2^+/WT^* crosses. Efficient recombination in endothelial cells was confirmed by PCR using DNA extracted from embryonic hearts (**Fig. 2A**).

**Figure 2.**
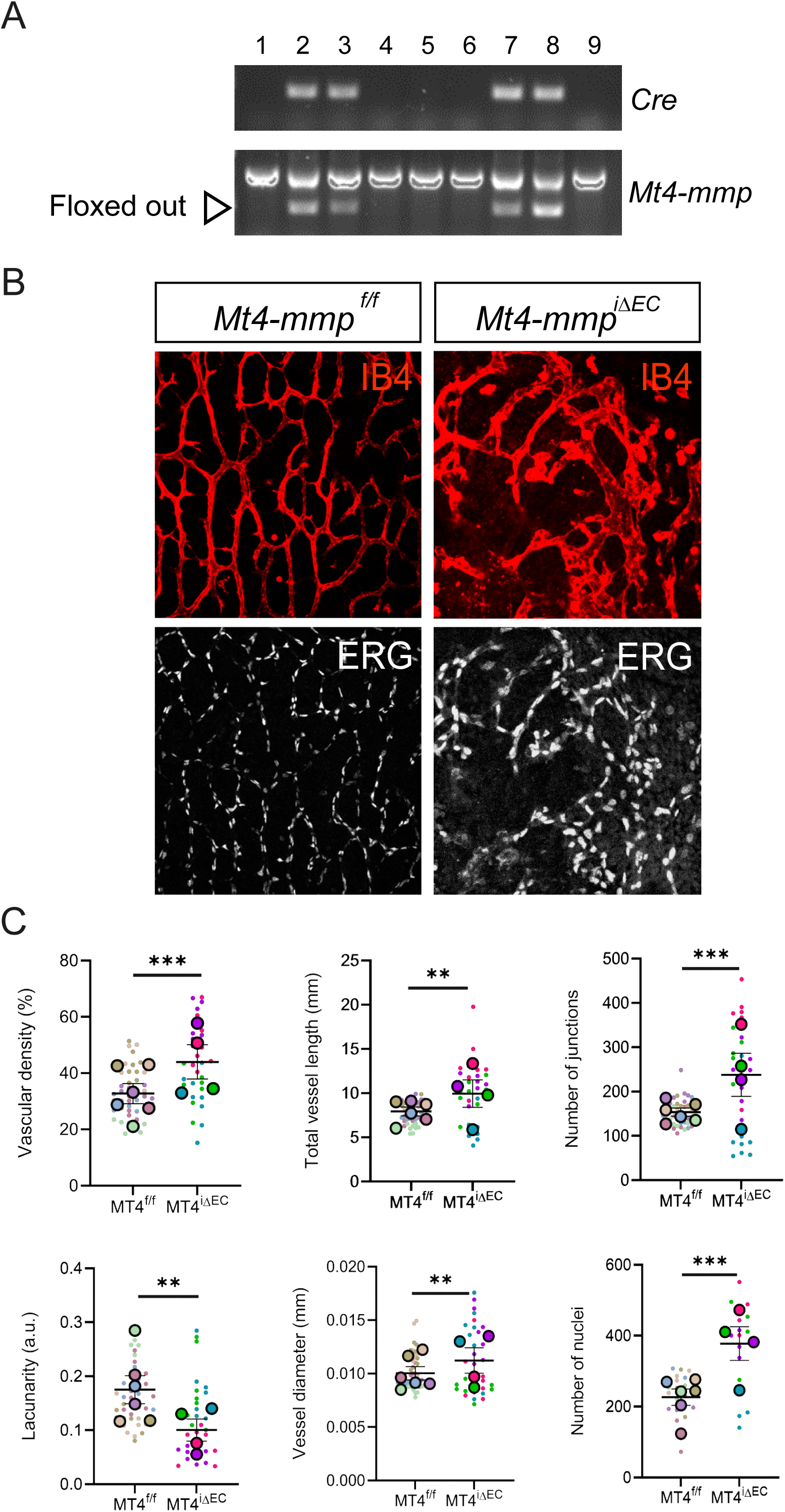
MT4-MMP deficiency in endothelial cells leads to enhanced angiogenesis in the E11.5 hindbrain. **A)** PCR analysis of the *Cre* and *Mt4-mmp* alleles in genomic DNA from E11.5 embryonic hearts obtained from pregnant mice from *Mt4-mmp^f/f^*;*Cdh5^CreERT2+/-^*crosses treated with tamoxifen from E7.5 to E9.5. **B)** Representative maximum intensity projection (MIP) images of whole-mount E11.5 hindbrains from *Mt4-mmp*^f/f^ and *Mt4-mmp*^iΔEC^ embryos, stained with IB4 (red); view shows the SVP. **C)** Quantification of vascular parameters (vascular density, total vessel length, number of junctions, and lacunarity) using AngioTool in the SVP of E11.5 hindbrains from *Mt4-mmp^f/f^*(n=6) and *Mt4-mmp^iΔEC^* (n=4) embryos. All the vascular parameters measurements refer to two different 0.25 mm^2^ crops per four hindbrain field acquired by confocal microscopy. Average vessel diameter was estimated as the ratio of vascular area to vessel length. Endothelial cell abundance was quantified as the number of ERG+ nuclei using ImageJ Values per individual image and the means ± SEM per embryo are shown in different colors. Data were analyzed using unpaired Welch’s t-test for parametric samples. p-value <0.01 **, <0.001 ***. Scale bar: 100 µm.

Analysis of IB4-stained E11.5 hindbrains revealed a striking opposite hypervascular phenotype compared to the global knockout. Loss of endothelial MT4-MMP led to exacerbated angiogenesis within the SVP, characterized by a highly disorganized and poorly patterned vascular network (**Fig. 2B)**. Quantitative analysis revealed a significant increase in vascular density, total vessel length and number of junctions, accompanied by reduced lacunarity in the SVP of *Mt4-mmp^i^*^ΔEC^ hindbrains compared to control *Mt4-mmp*^f/f^ embryos (**Fig. 2C**). This enhanced angiogenic phenotype was also associated with enlarged vessel diameter and increased numbers of endothelial cells (ERG-positive nuclei) (**Fig. 2C**). Therefore, loss of endothelial MT4-MMP compromised hindbrain angiogenesis in a distinct manner from global knockout at E11.5 (**Fig. 1**). This phenotypic divergence suggests a dual role for MT4-MMP: while the global absence of the protease likely impairs the overall angiogenic environment endothelial MT4-MMP specifically functions as a fine-tuner of vascular growth. In the absence of this endothelial-specific regulation, angiogenic expansion becomes unrestrained and disorganized.

Consistent with the global knockout, this phenotype was transient. By E12.5, *Mt4-mmp^i^*^ΔEC^ hindbrains displayed reduced vascular complexity, with fewer junctions, increased lacunarity, and decreased endothelial cell numbers, despite unchanged vessel diameters in the SVP (**Supp. Fig. S3A-B**). These findings suggest that endothelial MT4-MMP is required to fine-tune angiogenic growth and vascular organization within a narrow developmental window.

Beyond the embryonic hindbrain, MT4-MMP also exhibited context-dependent roles in angiogenesis during wound healing. In adult skin, *Mt4-mmp* expression (β-gal) was detected in most endothelial cells (ERG+) of the basal vascular plexus but were markedly downregulated during early wound healing (**Supp. Fig. S1C**). Thus, only a few or almost no β-gal–positive endothelial cells were detected in the granulation tissue and peripheral areas of the wound at days 3 and 6 (**Supp. Fig. S1C**). Notably, *Mt4-mmp*–null mice exhibited a significantly increased vascular density both within wounds and peripheral tissue during skin healing at day 6 post-injury compared with wild-type mice (**Supp. Fig. S2A**). This exacerbated angiogenic response correlated with an accelerated wound closure (**Supp. Fig. S2B**). Furthermore, quantitative proteomic analysis of skin collected at different time points after injury (days 0, 2, 4, 6, and 9 post-wound) revealed that the most notable changes in protein abundance occurred at day 6, with a significant enrichment in biological processes such as angiogenesis and others in *Mt4-mmp*^-/-^ mice compared to wild types (**Supp. Fig. S2C**). Indeed, mice with endothelial-specific *Mt4-mmp* deletion (after 5 days of tamoxifen administration to *Mt4-mmp^f/f^*;*Cdh5-CreERT2^+/WT^*or *Mt4-mmp^f/f^*;*Cdh5-CreERT2^WT/WT^* mice ^48^) recapitulated the exacerbated angiogenesis phenotype of *Mt4-mmp*^-/-^ mice (**Supp. Fig. S2D-E)**, showing enhanced vascular density in the wound and periphery 6 days post-injury compared to controls. *Mt4-mmp^i^*^ΔEC^ wounds also showed higher numbers of total and proliferative endothelial cells within and around the injury site, with no differences in healthy skin. This enhanced angiogenesis correlated with accelerated wound closure in *Mt4-mmp^i^*^ΔEC^ mice (**Supp Fig. S2F**).

Together, these results show that global and endothelial-specific loss of MT4-MMP differentially affect angiogenesis at E11.5, leading to reduced or increased vascularization, respectively. While global deficiency impairs vascular network formation, endothelial loss results in increased but disorganized vessel growth. Consistent with this, MT4-MMP deficiency also enhances angiogenesis during adult skin wound healing.

### MT4-MMP-deficient endothelial cells exhibit sustained VEGFA/ERK signaling and defective polarization

In both the embryonic hindbrain and adult skin wound healing, endothelial MT4-MMP deficiency disrupted angiogenic vessel directionality toward the brain ventricle or the wound bed, respectively, and increased endothelial cell numbers. To understand the molecular basis of this impaired vessel growth, we analyzed EC behavior *in vitro*. To this end, monolayers of mouse aortic endothelial cells (MAECs) from wild type and *Mt4-mmp*^-/-^ mice were cultured in the absence or presence of the angiogenic factor VEGFA and a scratch was inflicted to induce directional cell migration. Endothelial cell polarization was assessed by quantifying the angle between the nucleus-Golgi axis and a line perpendicular to the wound front (see M&M for details ^49^). As expected, most ECs (ECs) under all conditions display angles below 30°, indicating a proper polarization toward the wound edge. Notably, the average angle of MT4-MMP-deficient ECs in the first line (with or without VEGFA) and in the second line (with VEGFA) was significantly greater than in wild-type cells (**Fig. 3A**), indicating defective polarization, consistent with *in vivo* observations in skin wounds (**Supp. Fig. 2A, D**). Moreover, MT4-MMP-null endothelial cells exhibited faster migration than wild-type cells in the absence of VEGFA, with a similar trend observed upon VEGFA stimulation (**Fig. 3B**).VEGFA exerts its pro-angiogenic actions mainly by binding to VEGFR2, triggering its auto-phosphorylation and subsequent signaling that leads to ERK activation, which primarily drives endothelial cell proliferation but also migration, arterial fate specification and homeostasis ^50^. VEGFA can also activate other pathways, such as phospholipase PLCβ3, which regulates the directional migration of ECs ^51^. Indeed, VEGFA induced early phosphorylation of PLCβ3 (Ser537/Ser1105) in wild-type endothelial cells with a peak at 10 min and returning to baseline levels after 60 min of stimulation (**Fig. 3C**). MT4-MMP-null cells exhibited similar kinetics but elevated phospho-PLCβ3 levels (**Fig. 3C**). Moreover, while wild-type endothelial cells displayed a transient, biphasic activation of ERK1 (Thr202/Tyr204) and ERK2 (Thr185/Tyr187) in response to VEGFA (peaking at 5 and 30 min), MT4-MMP-deficient cells showed a sustained, plateau-like ERK activation that persisted up to 60 min (**Fig. 3C**). These signaling effects in the absence of MT4-MMP were selective, as no major changes were observed in VEGFA-induced p38 MAPK phosphorylation (Thr180/Tyr182) between wild-type and MT4-MMP-deficient endothelial cells (peaking at 10 min) (**Fig. 3C)**. Therefore, lack of MT4-MMP in ECs results in enhanced and sustained VEGFA-triggered signaling (ERK) that likely drives the increased EC proliferation and disorganized vessel patterning observed *in vivo*. Additionally, we observed elevated phosphorylation of PLCβ3, a key regulator of directional migration, but not in p38 MAPK involved in shear-stress-induced angiogenesis, permeability and survival ^50^ (**Fig. 3C**).

**Figure 3.**
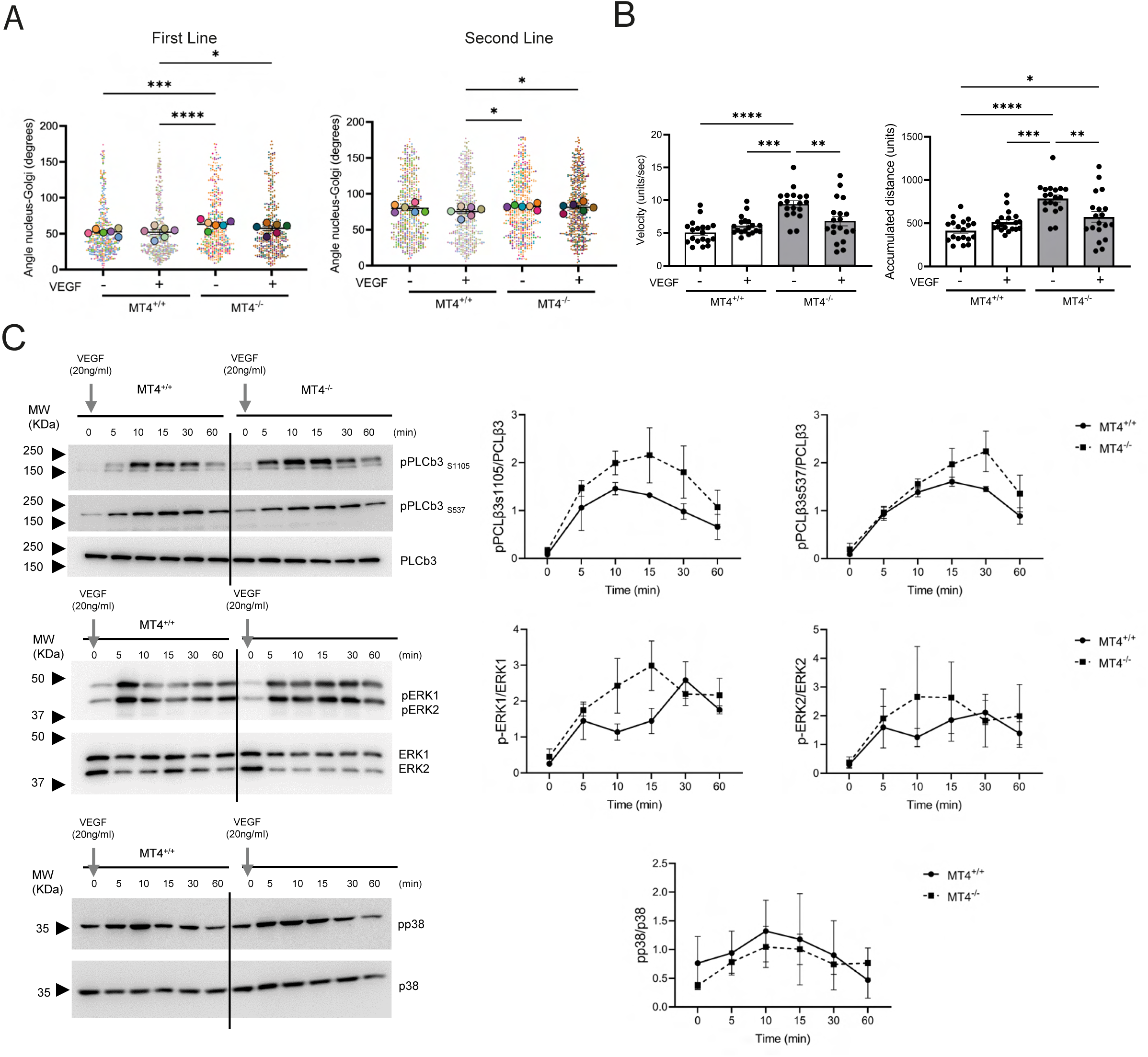
Absence of MT4-MMP results in altered endothelial cell polarization and sustained VEGF-mediated signaling *in vitro*. **A)** Polarization of *Mt4-mmp*^+/+^ and *Mt4-mmp*^-/-^ mouse aortic endothelial cells (MAECs) in scratch-wound assays. Graphs show the angle formed by the nucleus-Golgi axis relative to the wound front in the migrating cells at the first or second lines in the absence or presence of VEGFA (n > 30 cells per condition from 6 independent experiments). **B)** Quantification of migration speed and total distance measured in the migratory MAECs from *Mt4-mmp*^+/+^ and *Mt4-mmp*^-/-^ mice cultured and stimulated as in A. Individual cell values and the mean are shown (n ≥ 10) from 2 independent experiments. **C)** Monolayers of MAECs from *Mt4-mmp*^+/+^ and *Mt4-mmp*^-/-^ mice were serum-starved for 2 hours and stimulated with VEGFA (20 ng/ml) for different times (from 5 up to 60 min). Representative Western blots (left) and quantification (right) are shown for the analysis of phospho-PLCβ3 (S537 and S1105), phospho-ERK1 and ERK2 (Thr202/Tyr204; Thr185/Tyr187), and phospho-p38 MAPK (Thr180/Tyr182). n=3 independent experiments. Statistical analysis: Kruskall-Wallis test with Dunn’s posthoc (A, B) and a Two-way ANOVA (C). *p < 0.05; **p < 0.01; ***p < 0.001; ****p < 0.0001.

### NRP1 shedding is regulated by MT4-MMP in GPI-lipid domains

We next investigated how MT4-MMP deficiency modulates VEGFA-VEGFR2 signaling in ECs. We first hypothesized that MT4-MMP acts via proteolytic processing of a protein relevant to VEGFR2 signaling. We had previously identified novel MT4-MMP substrates using SILAC proteomics on the culture supernatant of primary bone marrow cell cultures established from mice expressing or not the protease ^23,52^. We screened the list of peptides for putative substrates (peptide_WT_/peptide_KO_ ratio ≥ 2) potentially contributing to phenotypes observed in MT4-MMP-null endothelial cells. Neuropilin-1 (NRP1) emerged as a compelling candidate (K.SFEGNNNYDTPELR.T peptide WT/KO ratio=1.96), given its role as a VEGFA co-receptor and a key regulator of angiogenesis ^53^. First, we confirmed the presence of a lower molecular weight NRP1 in the culture supernatants, with this shed form significantly more abundant in wild-type mouse bone marrow cells than in MT4-MMP-deficient cells, despite no significant differences in the total lysates (**Supp. Fig. S4A**). These findings indicate that MT4-MMP proteolytically sheds NRP1 from the cell surface. Next, we co-transfected HEK293 with mNRP1-EGFP and mouse MT4-MMP, either active or catalytically defective (E248A mutation ^23^). After confirming similar transfection efficiencies, we assessed NRP1 levels by anti-NRP1 staining and flow cytometry analysis in those HEK293 that co-expressed MT4-MMP and mNRP1-EGFP. We observed that NRP1 levels were higher on the cell surface of MT4-MMP/GFP-positive HEK293 cells expressing the catalytic mutant compared to those transfected with the MT4-MMP active form (**Supp. Fig. S4B**). These findings show that the proteolytic activity of MT4-MMP regulates NRP1 levels at the plasma membrane *in vitro*. Furthermore, NRP1 levels were significantly increased on the surface of endothelial cells from MT4-MMP-deficient mice obtained 6 days after skin injury from the periphery and the wound compared to those from wild-type mice (**Supp Fig. S4C**), indicating that NRP1 levels at the endothelial cell surface were regulated by MT4-MMP presence also *in vivo*.

Based on our established pipeline for identification of new substrates of MT-MMPs ^23,52,54^, CleavPredict analysis, protein modeling, and accessibility scoring identified the position 458 (S^458^Y) in NRP1’s b2 domain as the prime MT4-MMP cleavage site (**Fig. 4A**). *In vitro* digestion of a recombinant human NRP1 (Phe22-Lys852)-IgG1 Fc (Pro100-Lys330) chimera with the recombinant human catalytic domain of MT4-MMP yielded a ∼60 kDa C-terminal fragment (∼35 kDa post-IgG1 Fc subtraction), compatible with S^458^Y cleavage and proportional to protease concentration (**Fig. 4B**). Based on the presence of MT4-MMP in lipophilic fractions extracted with Triton X-114 ^52^, we observed that NRP1 was significantly more abundant in lipophilic fractions, but not in the hydrophilic fractions, of mouse endothelial cells lacking MT4-MMP compared to wild-type cells, under basal conditions and also after stimulation with VEGFA to a lesser extent (**Fig. 4C**). Interestingly, molecular dynamics simulations revealed that this proteolytic event is highly dependent on the lipid environment. The presence of phosphatidylserine (PS)-enriched membranes (up to 20% outer leaflet PS; typical of GPI-anchored protein domains) significantly stabilized the NRP1/MT4-MMP complex, shortening the distance between the protease catalytic pocket and the NRP1 cleavage site (**Fig. 4D; Supp Movies S1-S4**). Moreover, NRP1/MT4-MMP association was more stable in the plasma membrane with 20% PS as shown by the significantly lower energy of the complex (**Fig. 4D, Supp. Fig. S4D**). Therefore, the localization or recruitment of NRP1 to GPI or lipid domains could facilitate its cleavage by MT4-MMP, which is consistent with the fact that exposed PS regulates other protease-substrate pairs ^56^.

**Figure 4.**
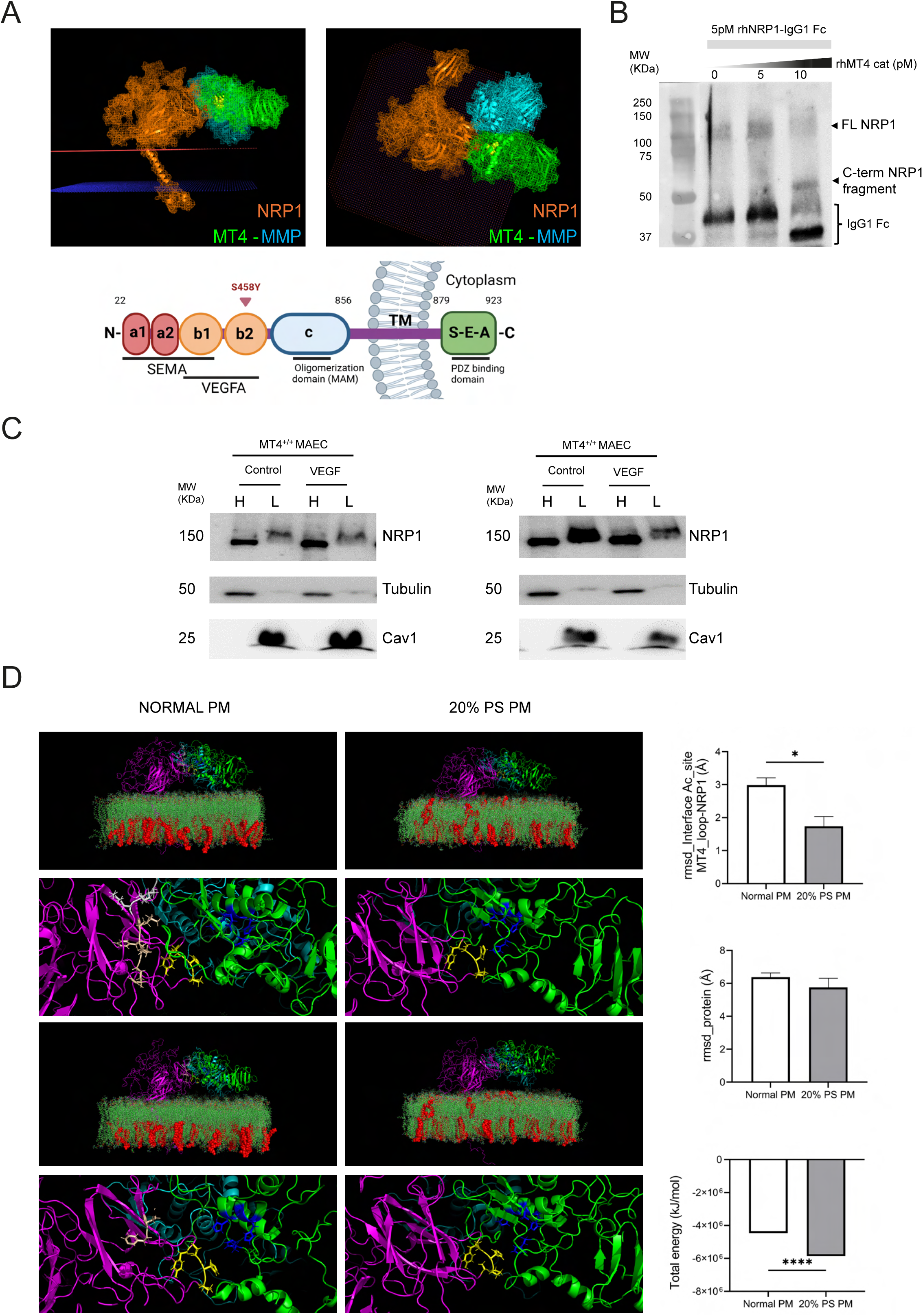
NRP1 is a novel substrate of MT4-MMP at lipid domains. **A)** *In silico* 3D modeling of the human NRP1 (orange) and MT4-MMP dimer complex (green and cyan), highlighting the putative S^458^Y cleavage site (red) in NPR1, the catalytic center of the protease (yellow) and the lipid bilayer (red & blue dots). 3D lateral (left) & top (right) views are shown. Depiction of human NRP1 domains, indicating the predicted cleavage site at position S^458^Y in the b2 domain (bottom, created in BioRender). **B)** *In vitro* digestion (n=5) of recombinant rhNRP1-IgG1 Fc (FL NRP1, 5 pM) with rhMT4-MMP catalytic domains (rhMT4 cat, 0,5 or 10 pM) developed with an anti-human IgG1 antibody to detect NRP1 C-terminal fragments. **C)** NRP1 levels assessed by Western blot of in Triton X114-extracted hydrophilic (H) and lipophilic (L) fractions from MAEC monolayers of *Mt4-mmp*^f/f^ and *Mt4-mmp*^iΔEC^ mice, serum-starved for 2 hours and stimulated with VEGFA (20 ng/ml) for 15 min. Tubulin and caveolin-1 are included as loading controls for hydrophilic and lipophilic fractions, respectively. A representative experiment of two with similar results is shown. **D)** *In silico* molecular dynamics simulations of human NRP1 (magenta) and MT4-MMP dimer (green and cyan) complex in plasma membrane with normal composition or containing 20% phosphatidylserine (PS, red) in the outer leaflet. Representative images of the complex and their magnified views are shown to the left at the initial (up) and final (down) steps of the simulation (see also **Movies S1-S4**). Graphs with the quantification of the distance between the NRP1 loop and MT4-MMP active site, between the proteins and the complex energy of the are shown to the right (n=3 independent replicas). Statistical analysis: Unpaired Student’s t-test. *p < 0.05; ****p < 0.0001.

We next explored whether NRP1 is accessible to MT4-MMP cleavage *in vivo* in embryonic hindbrain. We first performed *in situ* hybridization (ISH) for *Mt4-mmp* and *Nrp1* on sections of E11.5 hindbrains (**Fig. 5A**). *Mt4-mmp* mRNA expression was observed in regions typically occupied by neural progenitors and, albeit weakly, in a few ECs, primarily within vessels near the SVP (**Fig. 5A**). *Nrp1* mRNA levels were highly intense in neural progenitors and detected in scattered endothelial cells of the SVP (**Fig. 5A**). Complementarily, MT4-MMP and NRP1 immunostaining revealed that MT4-MMP is expressed primarily in a subset of vessels of the SVP proximal to the ventricle, but not in those of the PNVP, in the hindbrain of wild-type E11.5 embryos (**Fig. 5B**). NRP1, although faintly detectable, co-localized with MT4-MMP in some ECs of these wild-type hindbrains (**Fig. 5B**). As expected, the protease was not detected in sections of MT4-MMP-deficient embryonic hindbrains, where vascular NRP1 was also undetectable, likely reflecting the impaired angiogenic phenotype. We observed a similar pattern of MT4-MMP expression in vessels near the SVP and faint NRP1 staining in some of these vessels in hindbrains of *Mt4-mmp^f/f^* E11.5 embryos. Furthermore, we confirmed that MT4-MMP was absent from ECs in *Mt4-mmp^iΔEC^* hindbrains, whereas NRP1 expression persisted (**Fig. 5C**).

**Figure 5.**
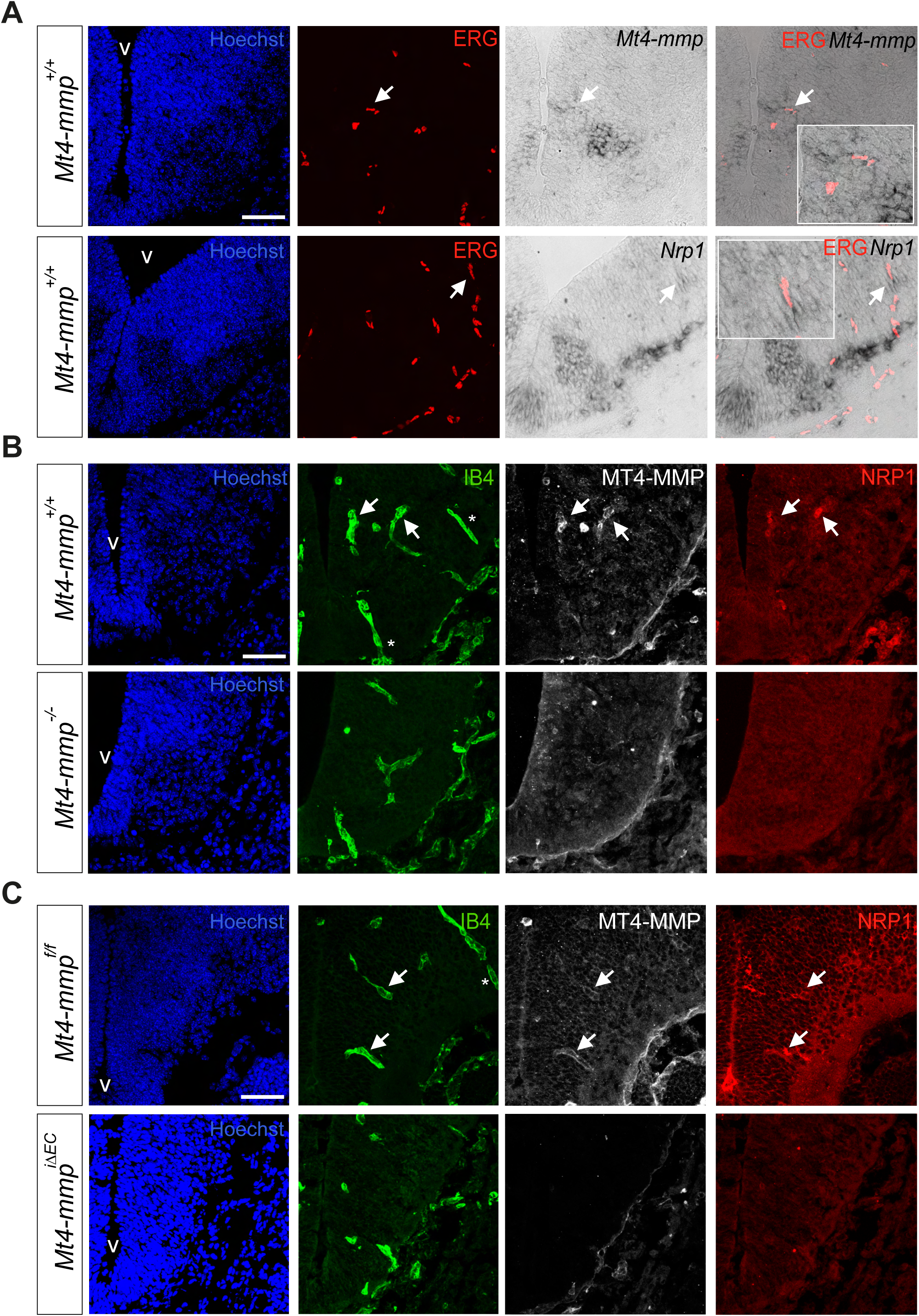
MT4-MMP and NRP1 are co-expressed in a subset of endothelial cells in the E11.5 developing hindbrain. **A)** Representative images of *in situ* hybridization (ISH) for *Mt4-mmp* and *Nrp1* and Maximum Intensity Projection (MIP) of ERG-stained endothelial cell nuclei in transverse sections of E11.5 wild-type hindbrains. Arrows indicate the expression of *Mt4-mmp* and *Nrp1* transcripts in endothelial cells. **B)** Representative MIP confocal images of E11.5 hindbrain sections of from *Mt4-mmp^+/+^* and *Mt4-mmp*^-/-^ embryos stained for IB4 (green), MT4-MMP (gray), NRP1 (red) and Hoechst (blue, nuclei). Arrows highlight endothelial cells near the subventricular plexus (SVP) exhibiting MT4-MMP and NRP1 co-localization. Asterisks indicate endothelial cells migrating from the perineural vascular plexus (PNVP) that lack detectable expression of both MT4-MMP and NRP1. **C)** Representative MIP confocal images from of E11.5 hindbrain sections from *Mt4-mmp*^f/f^ and *Mt4-mmp*^iΔEC^ embryos stained for IB4 (green), MT4-MMP (gray), NRP1 (red) and Hoechst (blue, nuclei). Arrows point to endothelial cells near the SVP showing co-localization of MT4-MMP and NRP1, whereas the asterisks mark endothelial cells penetrating from the PNVP that do not express either protein. Scale bar: 50 µm. V, ventricle.

These findings suggest that MT4-MMP acts as a spatial modulator rather than a constitutive protease, fine tuning NRP1 surface availability within specific membrane microdomains. By gating the intensity and duration of VEGFA/ERK signaling in SVP endothelial cells, MT4-MMP-mediated cleavage of NRP1 emerges as a key regulatory mechanism during hindbrain vascular development *in vivo*.

### MT4-MMP deficiency preserves VEGFA binding by modulating ERK signaling in the embryonic hindbrain

NRP1 binds VEGFA through its b1-b2 domains ^53^ (**Fig. 4A**). Given that the putative cleavage site S^458^Y is located at the C-terminal end of the b2 domain, and that MT4-MMP deficiency increases NRP1 surface levels *in vitro* (**Supp. Fig. S4B-C**), we assessed VEGFA binding capacity to hindbrain vessels using a VEGFA165 isoform fused to alkaline phosphatase (AP) ^26^. We applied this assay to embryonic hindbrains from mice lacking MT4-MMP either globally or selectively in endothelial cells. Although semi-quantitative ^26^, VEGFA binding (primarily to NRP1/VEGFR2 complexes) serves as a readout for NRP1 surface availability on blood vessels. VEGFA165-AP bound to the impaired vascular plexus of E11.5 *Mt4-mmp*^-/-^ hindbrains, exhibiting a discontinuous signal compared to wild types (**Supp. Fig S5A**). In conditional *Mt4-mmp^i^*^ΔEC^ mice, VEGFA165-AP bound to the vascular plexus of E11.5 hindbrains as in *Mt4-mmp^f/f^* controls (**Supp. Fig. 5B**), but in a pattern similarly to the abnormal vessels observed in IB4-stained hindbrains (**Fig. 2A-H**). These findings indicate that endothelial NRP1 retains the ability to bind to VEGFA in global and conditional MT4-MMP-null mice but possibly with a different efficiency.

Therefore, we assessed by immunostaining in hindbrain sections, VEGFA-induced signaling pathways potentially modulated by NRP1 *in vivo*, focusing on pERK, a downstream VEGFR2 target upregulated in cultured MT4-MMP-null ECs (**Fig. 3C**). Accordingly, pERK signal was consistently elevated in ECs of the vascular plexus of E11.5 hindbrains from *Mt4-mmp^i^*^ΔEC^ mice compared to *Mt4-mmp^f/f^* controls (**Fig. 6A-B**), correlating with the enhanced angiogenesis phenotype observed in these embryos (**Fig. 2**) and with the increased pERK observed in MAEC lacking MT4-MMP (**Fig. 3C**). In contrast, pERK signaling was reduced in ECs of E11.5 *Mt4-mmp*^-/-^ hindbrains compared to wild types, reflecting the reduced vascular plexus observed in these mice (**Fig 1**). In these mutant hindbrains, only some pERK-positive cells were detected in a few vessels coming from the PNVP, in which MT4-MMP was not expressed in wild-type mice (**Fig. 5B**).

**Figure 6.**
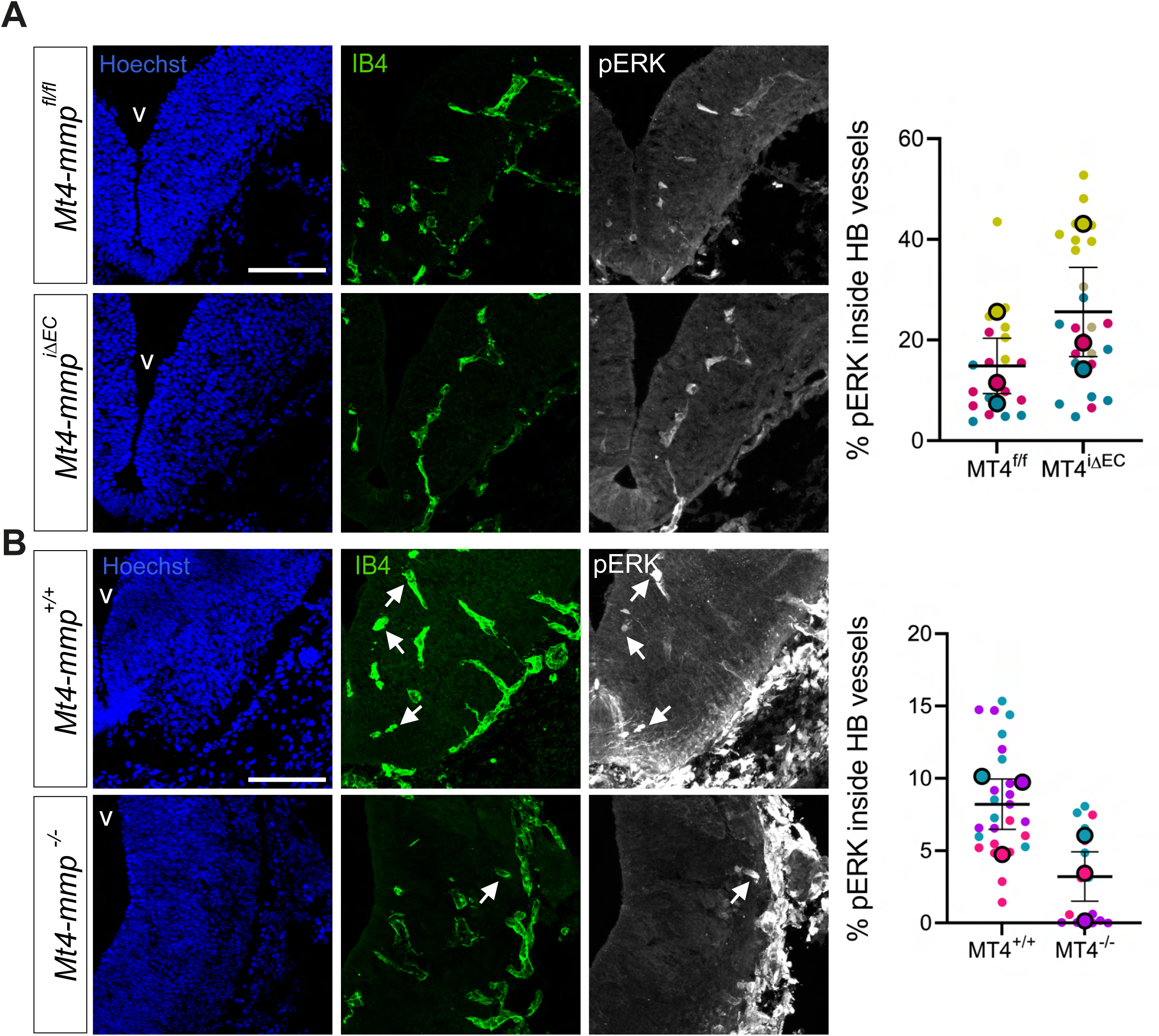
ERK signaling correlates with vascular phenotypes in embryonic hindbrains lacking MT4-MMP globally or selectively in endothelial cells. A, **B)** Representative MIP confocal images of E11.5 hindbrain sections from *Mt4-mmp*^f/f^ and *Mt4-mmp*^iΔEC^ embryos stained for IB4 (green), pERK (gray) and Hoechst (blue, nuclei), with corresponding quantification of pERK-positive signal within the vasculature (IB4+). **C, D)** Representative MIP confocal images of E11.5 hindbrain sections from *Mt4-mmp*^+/+^ and *Mt4-mmp*^-/-^ embryos stained for IB4 (green), MT4-MMP (gray), and Hoechst (blue, nuclei) and quantification of pERK positive signal within the vasculature (IB4+) of the hindbrain. For all quantifications, n=3 hindbrains per genotype were analyzed. Data points represent individual sections (colored dots) and the mean ± SEM per embryo (solid color). n=3 hindbrains for each genotype were analyzed, and values for each quantitated section and the mean ± SEM per embryo is shown in different colors. n=3 for each genotype. The comparison was performed using a paired Student’s t-test. Scale bar: 100 µm. V, ventricle.

These data indicate that ERK signaling is a primary effector of the MT4-MMP/NRP1 axis. Furthermore, they demonstrate that ERK activation is differentially regulated *in vivo* depending on whether MT4-MMP is lost only in endothelial cells or also in neighboring MT4-MMP/NRP1.

### Pharmacological inhibition of NRP1 partially rescues the vascular phenotype of MT4-MMP-deficient hindbrain

To confirm that the hypervascularization in *Mt4-mmp^i^*^ΔEC^ embryonic hindbrains stems from defective NRP1 regulation, we used the NRP1 inhibitor EG00229, which blocks VEGFA binding to NRP1 ^57^. Pregnant females were injected with tamoxifen i.p. from E7.5 to E9.5, followed by treatment with EG00229 or vehicle from E9.5 to E10.5, and embryos were analyzed at E11.5 (**Fig. 7A**). In vehicle-treated *Mt4-mmp^i^*^ΔEC^ embryos, the hindbrain vasculature showed significantly increased vessel density, reduced lacunarity, enlarged vessel diameters, and a trend toward increased vessel length and EC number of compared with vehicle-treated *Mt4-mmp^f/f^* controls (**Fig. 7A-B**), recapitulating most of the vascular phenotype observed without treatment (**Fig. 2**). Notably, pharmacological blockade of NRP1-VEGFA binding significantly rescued the vascular phenotype, reducing vascular density and restoring lacunarity to control levels (**Fig. 7A-B**). These results demonstrate that the MT4-MMP/NRP1 axis is a critical determinant of the vascular remodeling that prevents excessive and disorganized vessel growth during brain development acting specifically in ECs.

**Figure 7.**
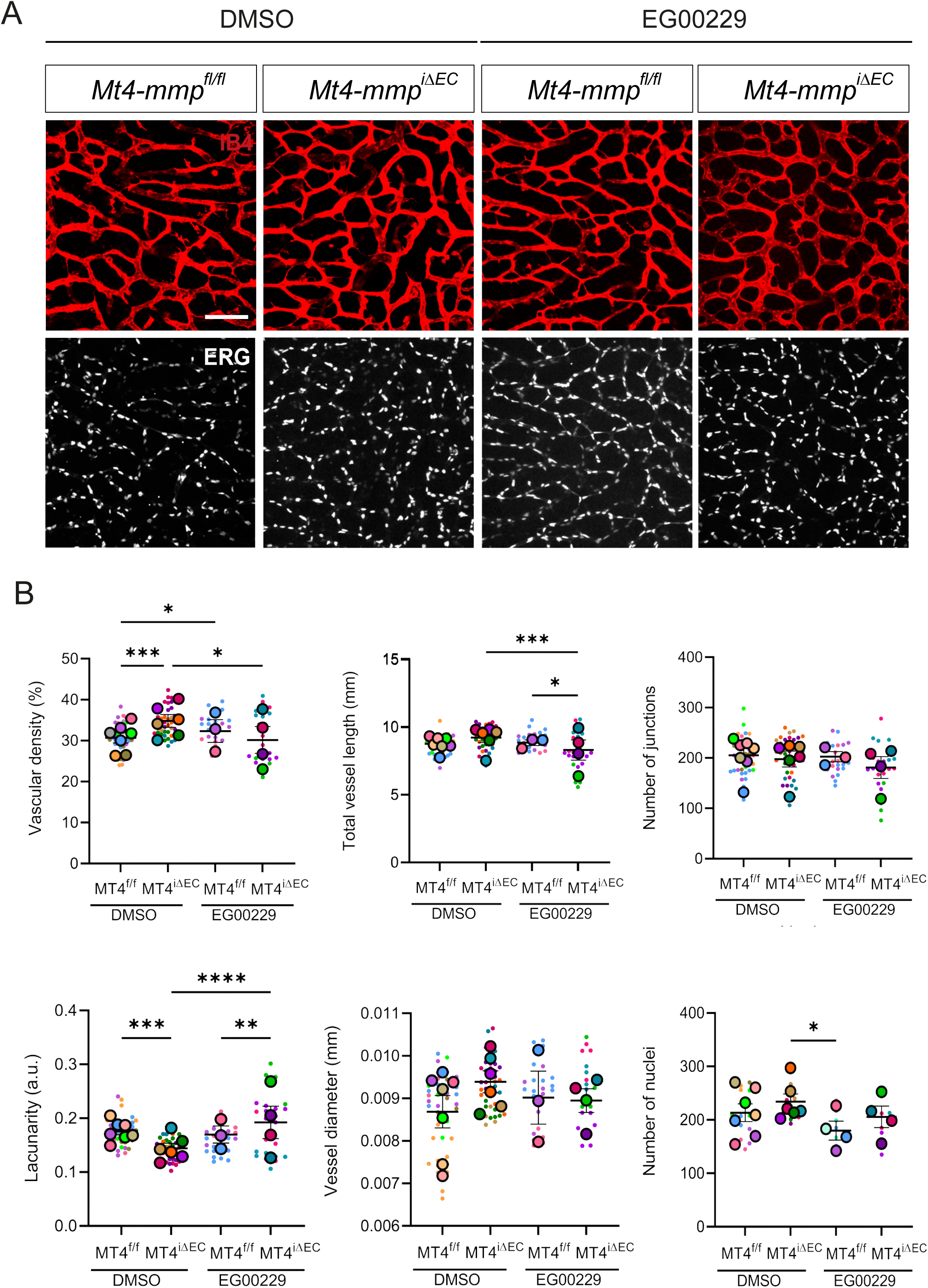
Inhibition of VEGF binding to NRP1 partially rescues the vascular phenotype in the hindbrains of E11.5 *Mt4-mmp*^iΔEC^ embryos. **A)** Representative MIP images of whole-mount E11.5 hindbrains from *Mt4-mmp*^f/f^ and *Mt4-mmp*^iΔEC^ embryos obtained from pregnant females treated with tamoxifen from E7.5 to E9.5 and with EG00229 or its corresponding vehicle (DMSO) at E9.5 and E10.5 and stained with IB4 (red) and anti-ERG (gray). **B)** Quantification of vascular parameters (vascular density, total vessel length, vascular junctions and lacunarity) with AngioTool in the SVP of *Mt4-mmp^f/f^* and *Mt4-mmp^iΔEC^* E11.5 embryos. Average vessel diameter was estimated as the ratio of vascular density and vessel length. Endothelial cell abundance was quantified using ImageJ by counting endothelial cell nuclei (ERG+). Data are shown as the individual image values and the mean ± SEM of (n=7 for *Mt4-mmp*^f/f^ with DMSO, n=6 for *Mt4-mmp*^iΔEC^ with DMSO, n=3 for *Mt4-mmp*^f/f^ with EG00229, and n= 4 for *Mt4-mmp*^iΔEC^ with EG00119). Statistical analysis: Kruskal-Wallis with Dunn’s post-hoc. p < 0.05; **p < 0.01; ***p < 0.001; ****p < 0.0001.

## DISCUSSION

Developmental angiogenesis is essential for neurovascular unit formation in the embryonic CNS, directing tightly regulated interactions among endothelial cells, neurons, and microglia. While the general vessel growth progression from the perineural vascular plexus (PNVP) to the subventricular vascular plexus (SVP) is well established in the developing brain, the molecular mechanisms ensuring its spatiotemporal organization remain elusive.

Here, using the embryonic mouse hindbrain as a model, we demonstrate that the MT4-MMP/NRP1/VEGFA/ERK signaling axis fine-tunes angiogenic progression within this dynamic neurovascular niche. These findings reveal a novel role for extracellular proteases in modulating VEGFA signaling during brain vascular development. Our study establishes the MT4-MMP/NRP1/VEGFA/ERK axis as a fundamental regulator of this process, where MT4-MMP acts as a context-dependent modulator that fine-tunes angiogenic growth across CNS development and adult wound healing.

### The MT4-MMP "Switch": Context-Dependent Regulation of Angiogenesis

It has previously been shown that MT4-MMP is involved in the regulation of vascular remodeling, particularly in pathological contexts such as tumor angiogenesis and in the aortic vessel wall development by acting primarily on perivascular cells ^23,58,59^. However, direct roles of MT4-MMP in endothelial cells have not yet been demonstrated. A major finding of this work is the striking phenotypic divergence between global and endothelial-specific MT4-MMP deficiency. While global loss impairs hindbrain vascularization, endothelial-specific deletion (*Mt4-mmp*^iΔEC^) triggers exacerbated, albeit disorganized, vessel growth. This contrast suggests that MT4-MMP coordinates a complex crosstalk between ECs and the surrounding neural environment.

The decreased vascular density and branching observed in MT4-MMP-null hindbrain angiogenesis resembled that of NRP1 mutants, either in the global knockout or the endothelial-selective deletion ^9,11^. In contrast, the increased vascular density and branching with more abundant endothelial cells in *Mt4-mmp*^iΔEC^ embryos resembled the actions of heparin-binding VEGFA isoforms such as VEGFA_165_ able to bind NRP1 ^7^. This was particularly inspiring, as the contrasting phenotypes observed between the global and endothelial cell-selective MT4-MMP-deficient embryos supports the notion that the crosstalk between ECs and surrounding cell types co-expressing MT4-MMP and NRP1 plays a crucial role. Indeed, our previous data in *Mt4-mmp*^LacZ/+^ reporter mice ^22^ and results in this report show that MT4-MMP and NRP1 are co-expressed in some ECs of the embryonic hindbrain but likely also in neural progenitors. Therefore, in the global MT4-MMP-deficient model, the absence of the protease could regulate NRP1 not only in endothelial cells but also in other cell types, leading to competition for factors such as VEGFA and/or trans-sequestering VEGFR2 from neighboring cells ^60^. In both scenarios, this would likely result in decreased VEGFR2 signaling in endothelial cells and, consequently, impaired angiogenesis. This was corroborated, to some extent, by the reduction in pERK signaling observed in endothelial cells of the SVP from *Mt4-mmp*^-/-^embryos. On the other hand, selective MT4-MMP deletion in ECs would affect NRP1 surface actions only in endothelial cells resulting in a phenotype that could resemble the overexpression of heparin-binding VEGFA isoforms ^7^. Sustained VEGFA signaling, particularly ERK but not p38 MAPK pathway, may result from increased VEGFR2 levels and/or delayed VEGFR2 internalization by association to highly expressed NRP1 at the cell surface of MT4-MMP-null endothelial cells ^14,60^. Our data also confirms that balanced ERK signaling in ECs is critical for developmental angiogenesis, likely through the regulation of the cell proliferation/cell cycle arrest balance, as has been described in other angiogenic contexts ^61,62^.

### NRP1 emerges as a novel MT4-MMP substrate that fine-tunes VEGFA signaling gradients during brain angiogenesis

MT4-MMP is an atypical protease with few recognized ECM substrates ^59^. In fact, we predicted a globular and compact conformation of the MT4-MMP dimer that would allow only small and flexible substrates, such as osteopontin ^23^ or transmembrane receptors such as integrin αM ^52^, to be accessible to the catalytic pocket. We identify NRP1, a multidomain transmembrane protein ^53^, as a novel substrate for MT4-MMP. Cleavage at S^458^Y position is predicted to release a1-a2, b1 and a portion of b2 domains, thereby disrupting SEMA3A and VEGFA binding to NRP1. Crucially, this cleavage site preserves the distal cell adhesion sequence (KHRENKVFMRKFKIGYSN) ^63^ and leaves the C-terminal stalk intact. This potentially allows NRP1 to maintain its interaction with integrin αVβ8 to regulate TGFβ signaling ^20^. Furthermore, retention of the transmembrane and cytosolic domains may continue to drive intracellular signaling or regulate integrin trafficking, suggesting that MT4-MMP-mediated processing functions as a molecular switch that selectively terminates ligand-dependent signaling while preserving other co-receptor activities^53^.

Since VEGFA binds to the NRP1b1/b2 domains, MT4-MMP–mediated cleavage likely acts as a molecular switch, restricting VEGFA binding in wild-type ECs while enhancing it in MT4-MMP-deficient ECs. This model is supported by the observed opposing vascular phenotypes and the partial rescue of the exacerbated angiogenesis in *Mt4-mmp^i^*^ΔEC^ embryos using the inhibitor of VEGFA binding to NRP1, EG002229. The mechanism identified herein adds a critical layer of regulation to NRP1-dependent vascular actions, contributing to the generation of a NRP1/VEGFA signaling gradient during developmental angiogenesis. Previous reports on the absence of vascular phenotypes in NRP1 knock-in mouse models carrying point mutations that abolish VEGFA binding ^14^ suggest that functional signaling gradients may be especially relevant and only achieved if fine-tuned regulation occurs as we propose with MT4-MMP-mediated cleavage. Cleavage by MT4-MMP might also enable or unmask NRP1 binding sites for unidentified ligands, as previously hypothesized ^14^. In this line, we previously reported that cleavage of matricellular proteins - such as osteopontin (OPN) and thrombospondin (TSP) by MT4-MMP and MT1-MMP, respectively - enables integrin binding leading to JNK activation in smooth muscle cells ^23^ or nitric oxide production by ECs ^48^. Our observations in mouse models suggest that the absence of proteolytic processing often combines features of full-length substrate overexpression with a deficiency in specific interactions driven by cleavage-unmasked adhesion sites, as demonstrated for MT4-MMP/OPN in the aortic wall ^23^. Consequently, the vascular phenotype of *Mt4-mmp^i^*^ΔEC^ mice could combine features of both sustained VEGFA/VEGFR2 signaling (manifesting as increased vascular density ^7,64^) and dysregulated adhesion-driven signals or impaired TGFβ regulation (resulting in dilated vessels and failed anastomosis ^20^).

Although soluble forms of NRP1 are generated by alternative splicing ^53^, proteolytic shedding of NRP1 by transmembrane proteases had not been described until a recent report showing that ADAM9 and ADAM10 cleave NRP1 ^65^. NRP1 cleavage site was close to the membrane as expected for ADAM proteases, resulting in the generation of a C-terminal NRP1 fragment that inhibited VEGFA signaling ^65^. The impact that this cleavage may have on brain vascularization remained to be investigated. Furthermore, given that NRP1 may be present in lipid rafts where it is regulated by SEMA3A ^66^, MT4-MMP cleavage would only affect the NRP1 pool in these membrane domains. Our computational modeling supports this idea and adds a sophisticated layer to this mechanism for a more precise regulation in angiogenesis. The stabilization of the MT4-MMP/NRP1 complex in phosphatidylserine (PS)-enriched microdomains suggests that MT4-MMP does not act as a constitutive scavenger, but as a spatially restricted regulator. This provides a precise mechanism to generate local NRP1/VEGFA signaling gradients, which are essential for directional vessel ingression.

Unlike the VEGF-induced ADAM9/10-mediated shedding described by Mehta *et al.* ^65^, MT4-MMP-mediated cleavage appears to be restricted to specific lipid domains. This spatial compartmentalization suggests that MT4-MMP does not target the total surface NRP1, but rather a specific membrane pool, potentially modulating the availability of NRP1 for VEGFA signaling by regulating its distribution across different membrane environments.

Although the cleavage site was identified through computational modeling and *in vitro* digestion of the human NRP1 protein, SILAC-based proteomic analysis was performed in mouse cells. Furthermore, we observed consistent phenotypes both *in vitro* and *in vivo* in mouse models, suggesting that the orthologous cleavage site mediates the actions of NRP1 in mice. In this regard, the human and murine NRP1 sequences surrounding S^458^Y share 62,5% identity. More significantly, the proline at position P3’ is conserved in both human and murine NRP1; this residue confers the highest predicted score in the MT4-MMP cleavage matrix (MEROPS database, https://www.ebi.ac.uk/merops/cgi-bin/pepsum?id=M10.017;type=peptidase). Interestingly, our findings complement the recent discovery of MT6-MMP (MMP25) in brain angiogenesis ^67^. We find it interesting that the two unique GPI-anchored MT-MMPs are responsible for critical steps during cerebral angiogenesis: MT6-MMP facilitates the initial invasion of ECs into the pial basement membrane (through the processing of collagen IV), while MT4-MMP ensures the subsequent organized vessel branching reaching the ventricle through the processing of NRP1.

### Clinical Implications: From Wound Healing to Vascular Malformations

The actions of MT4-MMP on NRP1 signaling likely extend to other angiogenic scenarios beyond CNS development. In the context of skin healing, the effect appears to depend primarily on endothelial MT4-MMP, as evidenced by the similarity in vascular phenotype between global and endothelial-specific deficiency models. Furthermore, MT4-MMP deletion in smooth muscle cells or hematopoietic cells failed to recapitulate the phenotype observed in global deficiency mice (data not shown). MT4-MMP is expressed in basal endothelial cells of the skin and is downregulated in angiogenic vessels, contrary to what would be expected for NRP1 ^68^, which could create an NRP1 ligand binding gradient that is disrupted when MT4-MMP is absent from the basal skin in knockout mouse models. Moreover, the vascular phenotype correlated with accelerated wound closure, indicating that the MT4-MMP/NRP1 axis could serve as a target for promoting tissue healing. These findings would reinforce the concept of reactivation of embryonic programs in pathophysiological contexts and could open new therapeutic opportunities for repair after damage, not only in the skin but also in other tissues.

MT4-MMP/NRP1/ERK axis may also be of relevance in vascular malformations. Somatic mutations in the KRAS proto-oncogene resulting in activation of downstream ERK has been described in sporadic arteriovenous malformations (AVM), with inhibition of ERK pathway, but not PI3K/Akt, reversing the altered endothelial cell phenotype *in vitro* ^69^. Similarly, mutations in Gαq gene enhance PLCβ3 and ERK signaling resulting in capillary malformations in congenital Sturge-Weber Syndrome ^70^. Although no NRP1 mutations have been described yet in AVM patients, given the hyperactivation of ERK in endothelial cells lacking MT4-MMP, it is still possible that mutations affecting MT4-MMP catalytic activity may be associated with vascular malformations. In this regard, we previously identified MT4-MMP mutations in human patients affected by aortic aneurysms and demonstrated that protease absence resulted in higher predisposition to aortic aneurysms in mouse models ^23^. Therefore, the MT4-MMP/NRP1/ERK axis represents a novel pathway that could be targeted for the treatment of vascular malformations.

Finally, this regulatory axis may be relevant to SARS-CoV-2 infection, given that NRP1 binds the viral spike (S) protein. MT4-MMP cleavage at S^458^Y could prevent spike binding to NRP1, potentially protecting target cells; this stands in contrast to MT1-MMP, which promotes coronavirus entry and infection ^71^. Such divergent roles underscore the need for specific inhibitors and cell type-selective strategies when considering MT-MMPs as potential therapeutic targets in infectious and vascular diseases.

In summary, we have uncovered the MT4-MMP/NRP1/VEGFA/ERK axis as fundamental in neurovascular development, establishing pericellular proteolysis as a critical regulator for endothelial responses within the embryonic brain niche. Our findings reveal that endothelial protease MT4-MMP functions as a spatiotemporal controller on NRP1-dependent VEGFA signaling, preventing aberrant angiogenesis while ensuring proper vessel patterning. Through integrated proteomics, structural modeling, and functional assays, we establish NRP1 as a novel MT4-MMP substrate whose membrane-restricted cleavage fine-tunes VEGFA/ERK signaling to balance endothelial proliferation versus directional migration. This mechanism extends beyond development to govern adult skin wound repair, where MT4-MMP deficiency accelerates angiogenesis and wound closure. These findings establish MT4-MMP/NRP1 as a key regulator of vascular homeostasis with broad implications for neurovascular disorders, ischemic injury, and pathological angiogenesis.

## Supporting information

S2 Movie (zoom)

S4 Movie (zoom)

## Acknowledgements

We would like to thank Dr. Motoharu Seiki (Tokyo, Japan) for kindly providing us with the *Mt4-mmp*^lacZ/+^ reporter mice, Dr. Francesc Canals (Vall d’Hebron Institute of Oncology, Barcelona) for performing SILAC assays in the past and Dr. Alessandro Fantin (University of Milano) for kindly providing us with AP-VEGF ligands used in this study. Finally, we thank the Microscopy & Animal Facility Units at the Center for Biological Research Margarita Salas (CIB-CSIC) for technical assistance in this project.

## SUPPLEMENTARY FIGURES

**Supp. Figure S1. The protease MT4-MMP is expressed in endothelial cells of the mouse embryonic CNS and adult skin. A)** β-gal staining of E10.5 and E11.5 transverse sections from Mt4-mmp^LacZ/+^ embryos highlights *Mt4-mmp* transcriptional activation in blood vessels of the PNVP and those extending toward the ventricle (arrows; n=8). Detail of LacZ staining in the PNVP (arrows) and vessels within the brain parenchyma (open arrows). **B)** Immunostaining of E11.5 spinal cord transverse sections for the endothelial marker CD31 (green) and β-gal (red). Arrows indicate endothelial cells expressing β-gal in the blood vessels in the neural tube and the PNVP (open arrows). Abbreviations: MN, motoneurons; PNVN, perineural vascular plexus. **C)** Representative maximum intensity projection images of sections of the skin (basal, wound and periphery of the wound after 3 or 6 days) from Mt4-mmp^LacZ/+^ mice stained for β-gal (red), the endothelial markers CD31 (green) and ERG (gray), and Hoechst (blue, nuclei). Merged images are shown to the bottom. Scale bars: 100 µm (A: a, b, d), 50 µm (A: e) and 25 µm (A: c, f); and 100 µm (B, C).

**Supp Figure S2. Enhanced angiogenesis and accelerated closure during skin wound-healing in mice lacking MT4-MMP globally or selectively in endothelial cells. A)** Maximum intensity projection (MIP) images of wounded skin of *Mt4-mmp*^+/+^ (n=9) and *Mt4-mmp*^-/-^ (n=8) mice after 6 and 9 days stained in whole-mount with anti-CD31 (red). Quantification of vascular density (in the wound and the wound periphery) at different time-points after wound is shown to the right. Dotted lines mark the the wound border (W). **B)** Wound closure kinetics in *Mt4-mmp*^+/+^ and *Mt4-mmp*^-/-^ mice (n=13 mice per genotype). **C)** Functional categories significantly altered in wounded skin from *Mt4-mmp*^-/-^ versus *Mt4-mmp*^+/+^ mice after 6 days. Zc represents standardized log_2_ category ratios: negative values denote significantly decreased categories, whereas positive values indicate significantly increased categories. Circle size corresponds to the number of proteins included in each category, as indicated in the legend. **D)** Representative MIP images of the vasculature of the wounded skin from *Mt4-mmp*^f/f^ (n=9) and *Mt4-mmp*^iΔEC^ (n=8) mice stained with anti-CD31 (red) in whole-mount preparations. Quantification of the vascular density in the wound and the wound periphery. **E)** Representative confocal images (left) of the wounded skin of *Mt4-mmp^f/f^* and *Mt4-mmp^iΔEC^*(n=6 for each genotype) at day 6 stained for ERG (gray) and Ki67 (red). Quantification of the percentage of endothelial cells (ERG+) present in basal (unwounded) and wounded skin (W, wound; P, periphery), as well as and the percentage of proliferative endothelial cells (ERG+Ki67+) in the same areas, is shown to the right. **F)** Wound-closure kinetics in *Mt4-mmp*^f/f^ (n=7) and *Mt4-mmp*^iΔEC^ (n=5) mice. Data are shown as individual values, the means ± SEM (D, E) or as the means ± SEM (A, B, F). Statistical analysis was performed using an unpaired Student’s t test (D, E) or two-way ANOVA with Šídák’s multiple-comparisons test (B, F). p<-. *p < 0.05; **p < 0.01; ***p < 0.001; ****p < 0.0001. Scale bar: 100 µm.

**Supp. Figure S3. The vascular phenotype in hindbrains of *Mt4-mmp*^iΔEC^ embryos is transient. A)** Representative maximum intensity projection (MIP) images of embryonic hindbrains from *Mt4-mmp*^f/f^ and *Mt4-mmp*^iΔEC^ mice stained with IB4 (red) and anti-ERG (gray) in whole-mount preparations at E12.5; view from the SVP. **B)** Quantification with AngioTool of vascular parameters (vascular density, total vessel length, vascular junctions and lacunarity) in the SVP of E12.5 hindbrains from *Mt4-mmp^f/f^* (n=4) and *Mt4-mmp^iΔEC^* (n=3) embryos. All the vascular parameters measurements refer to two different 0.25 mm^2^ ROI per four hindbrain fields acquired in the confocal microscope. Average vessel diameter was estimated by vascular area/vessel length and the abundance of endothelial cells quantitated as ERG+ nuclei with ImageJ. Values per individual image and the means ± SEM per embryo are shown in different colors. Statistical comparisons were performed using Welch’s unpaired t test.. *p < 0.05; **p < 0.01; ***p < 0.001; ****p < 0.0001. Scale bar: 100 µm.

**Supp. Figure S4. Neuropilin-1 is a new substrate of the protease MT4-MMP. A)** Western blot analysis of NRP1 in culture supernatants and lysates of bone marrow-derived cells from *Mt4-mmp*^+/+^ and *Mt4-mmp*^-/-^ mice (left) and quantification of NRP1 levels normalized to Ponceau and tubulin as loading controls for the supernatant and lysate, respectively (right). n=2 samples per genotype. **B)** NRP1 cell surface levels assessed by flow cytometry in HEK293 cells co-transfected with mNRP1-EGFP and MT4-MMP, either wild-type or catalytically dead mutant (E248A), at 3:1 (protease:NRP1) ratio. Transfection efficiency was similar for all plasmids (not shown). On the left, a representative histogram of anti-NRP1 expression assessed in HEK293 cells co-expressing MT4-MMP and mNRP1-EGFP (empty line for wild-type MT4-MMP and solid gray line for E248A MT4-MMP), and the quantification of the geometric mean fluorescence intensity (Geo MFI) and median fluorescence intensity (Median MFI) of NRP1 on the right (n=3 independent experiments). **C)** Flow cytometry analysis of surface NRP1 levels in endothelial cells isolated from wound and wound periphery at day 6 in *Mt4-mmp*^f/f^ and *Mt4-mmp*^iΔEC^ mice (n=6 mice per genotype). Graphs show the normalized mean fluorescence intensity (MFI).. **D)** Energy profiles of the NRP1/MT4-MMP complex during molecular dynamics simulation in membranes of normal composition (left) or with 20% PS on the outer leaflet (right), estimated using GROMACS. Statistical analyses: paired t-test (A) and unpaired t-test (B, C). p < 0.05; **p < 0.01.

**Supp. Figure S5. VEGFA binding to NRP1 persists in hindbrains lacking MT4-MMP globally or selectively in endothelial cells. A)** Representative whole-mount images of AP control or VEGFA165-AP binding in E11.5 hindbrains from *Mt4-mmp*^+/+^ (n=4) and *Mt4-mmp*^-/-^ (n=9) embryos. **B)** Representative VEGFA165-AP and AP control binding images in E11.5 embryonic hindbrains from *Mt4-mmp*^f/f^ (n=3) and *Mt4-mmp*^iΔEC^ (n=5). Scale bar: 100 µm

**Supp. Movies S1 and S2. *In silico* molecular dynamics simulation of NRP1/MT4-MMP complex in a plasma membrane with normal composition. Movie S1:** Representative molecular dynamics simulation of the complex formed by human NRP1 (magenta) and the MT4 MMP dimer (green and cyan) embedded in a plasma membrane with normal lipid composition. **Movie S2:** Magnified view of the same simulation highlighting local interactions within the NRP1 and MT4 MMP interface.

**Supp. Movies S3 and S4. *In silico* molecular dynamics simulation of NRP1/MT4-MMP complex in a plasma membrane exposing phosphatidylserine in the outer leaflet. Movie S3:** Representative molecular dynamics simulation of human NRP1 (magenta) bound to the MT4 MMP dimer (green and cyan) in a plasma membrane containing 20% phosphatidylserine (red) in the outer leaflet. **Movie S4:** Magnified view of the NRP1 and MT4-MMP interface under the same phosphatidylserine enriched membrane conditions.

