## Supplementary figures and images for "MT4-MMP-mediated NRP1 shedding fine-tunes VEGFA signaling dynamics during embryonic brain angiogenesis"

### Supplementary Figures

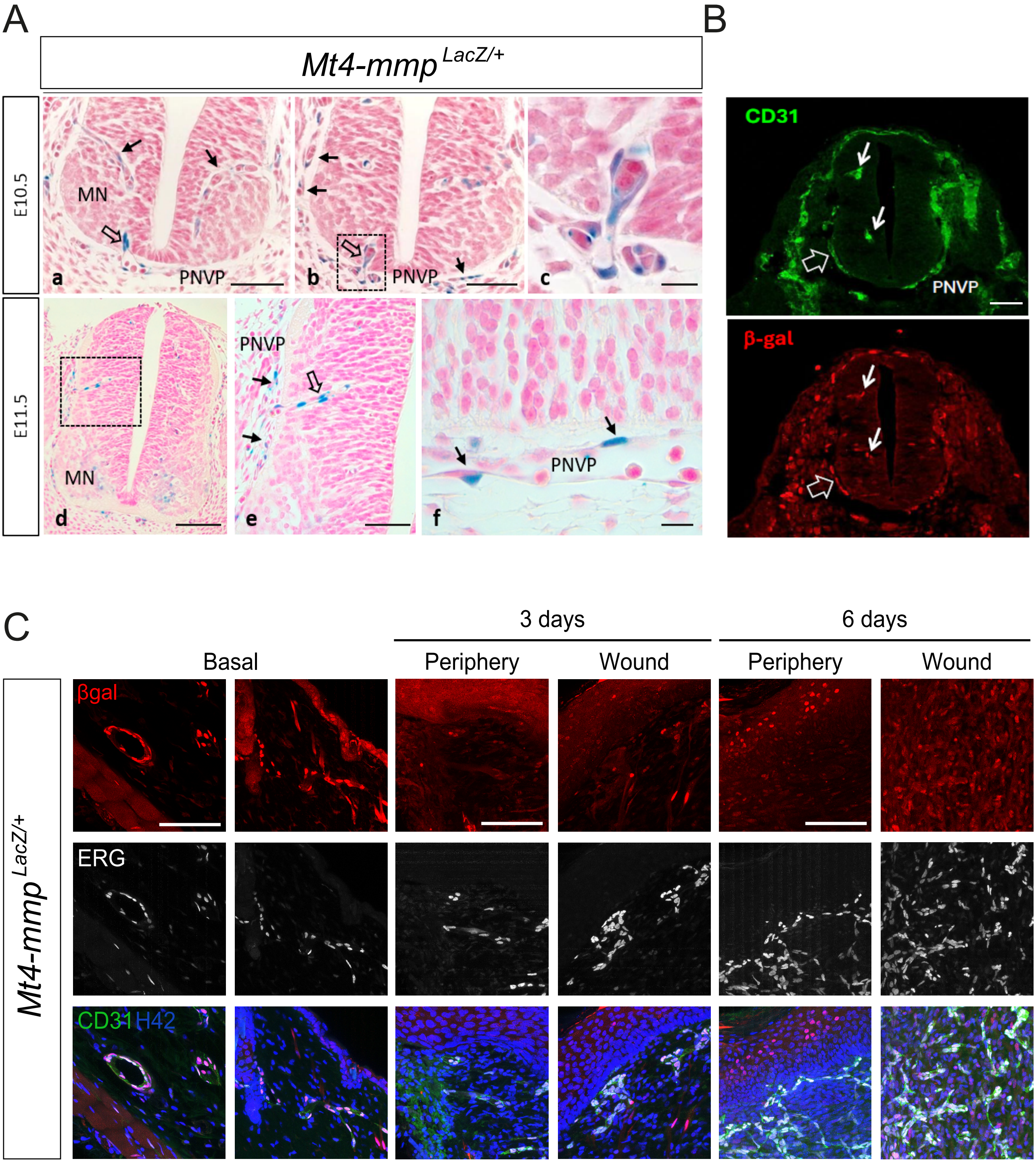

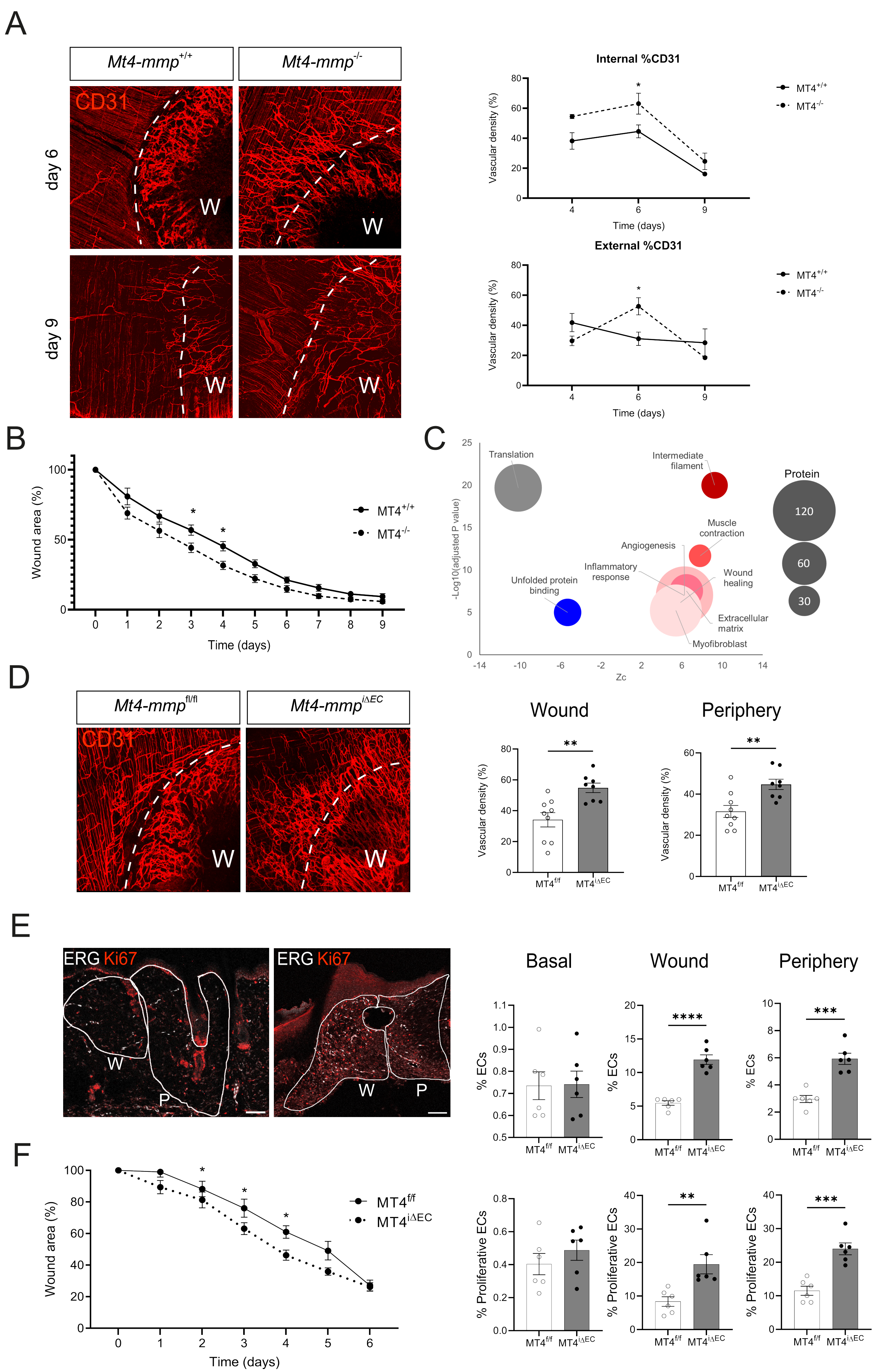

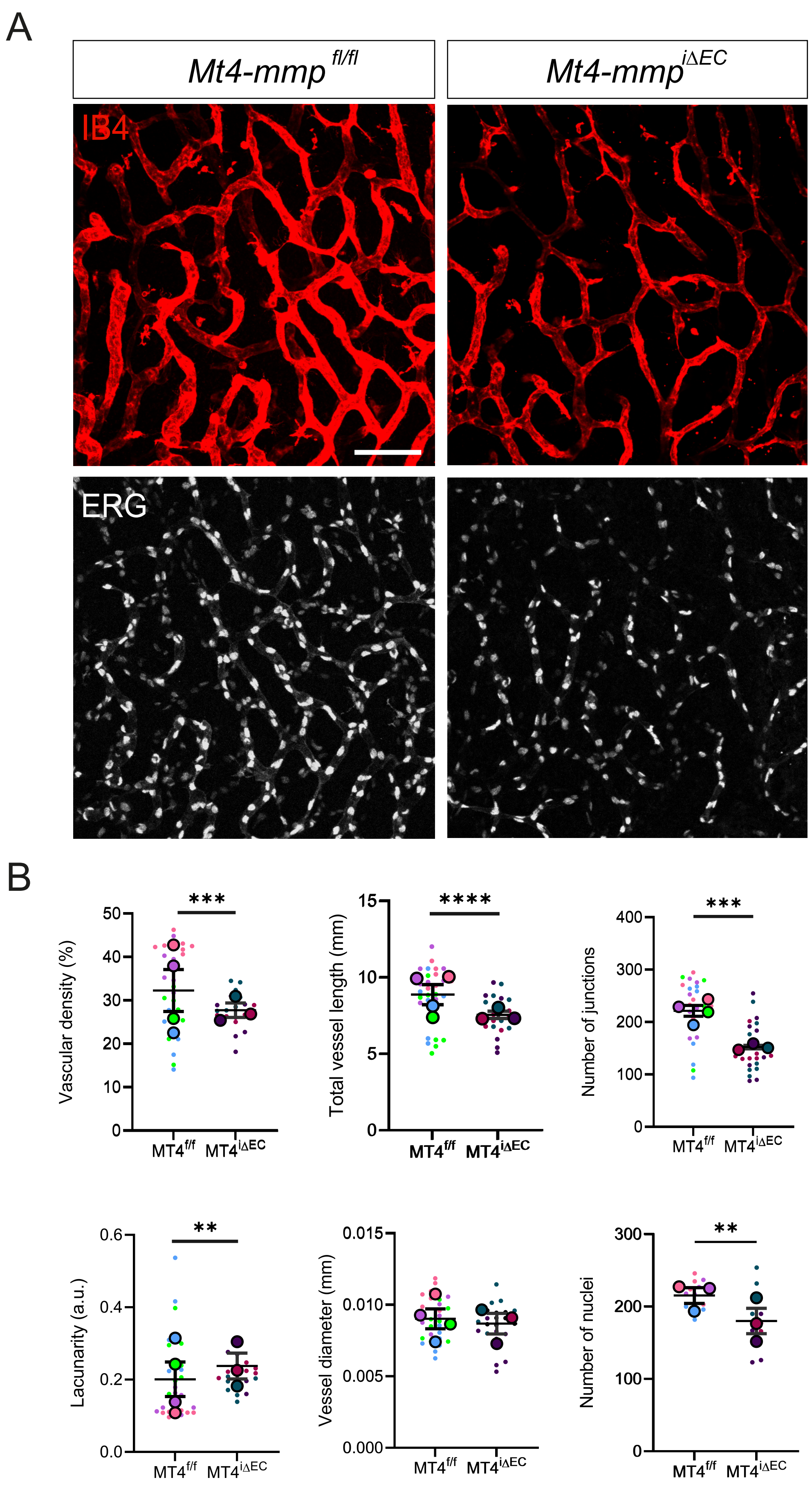

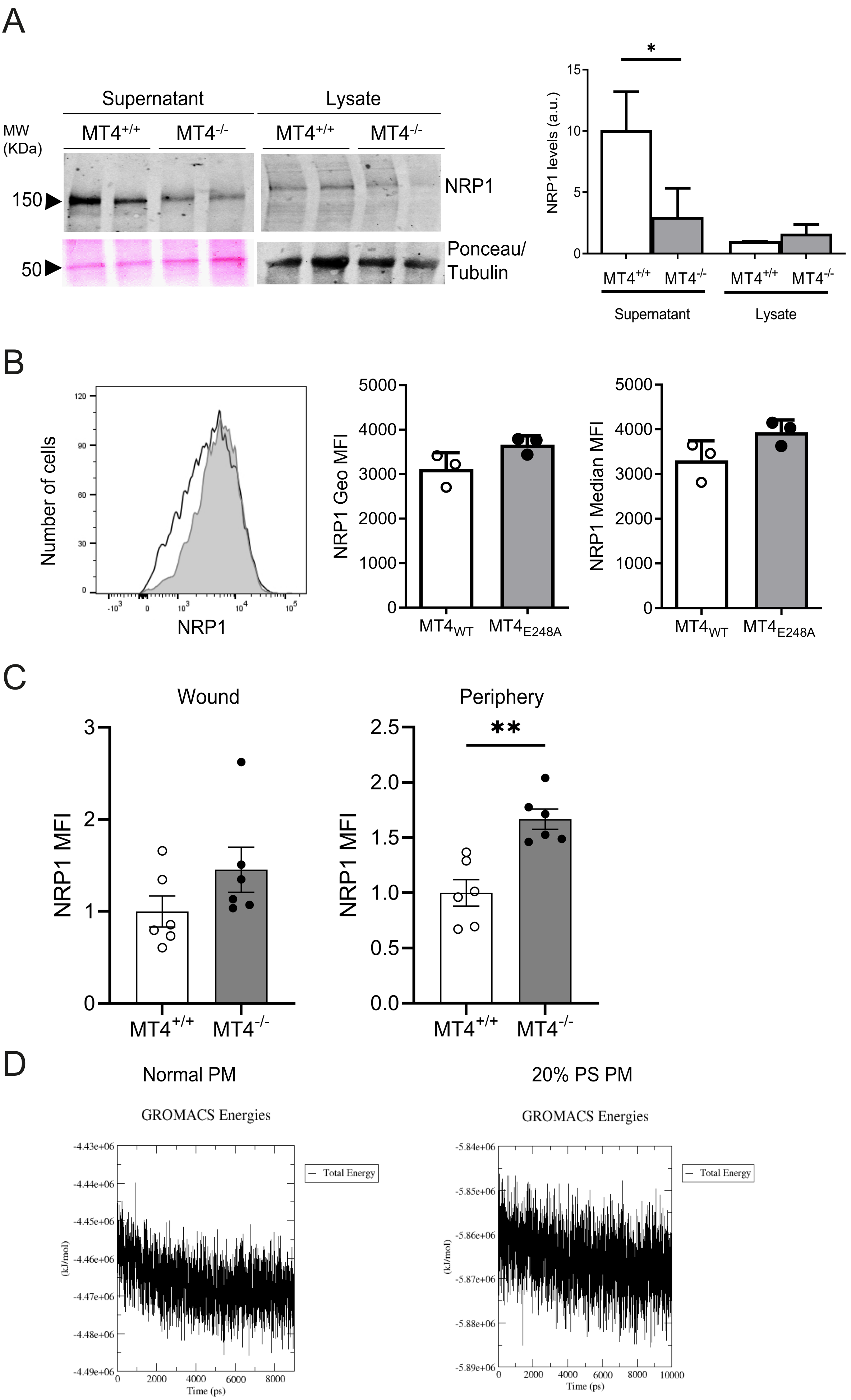

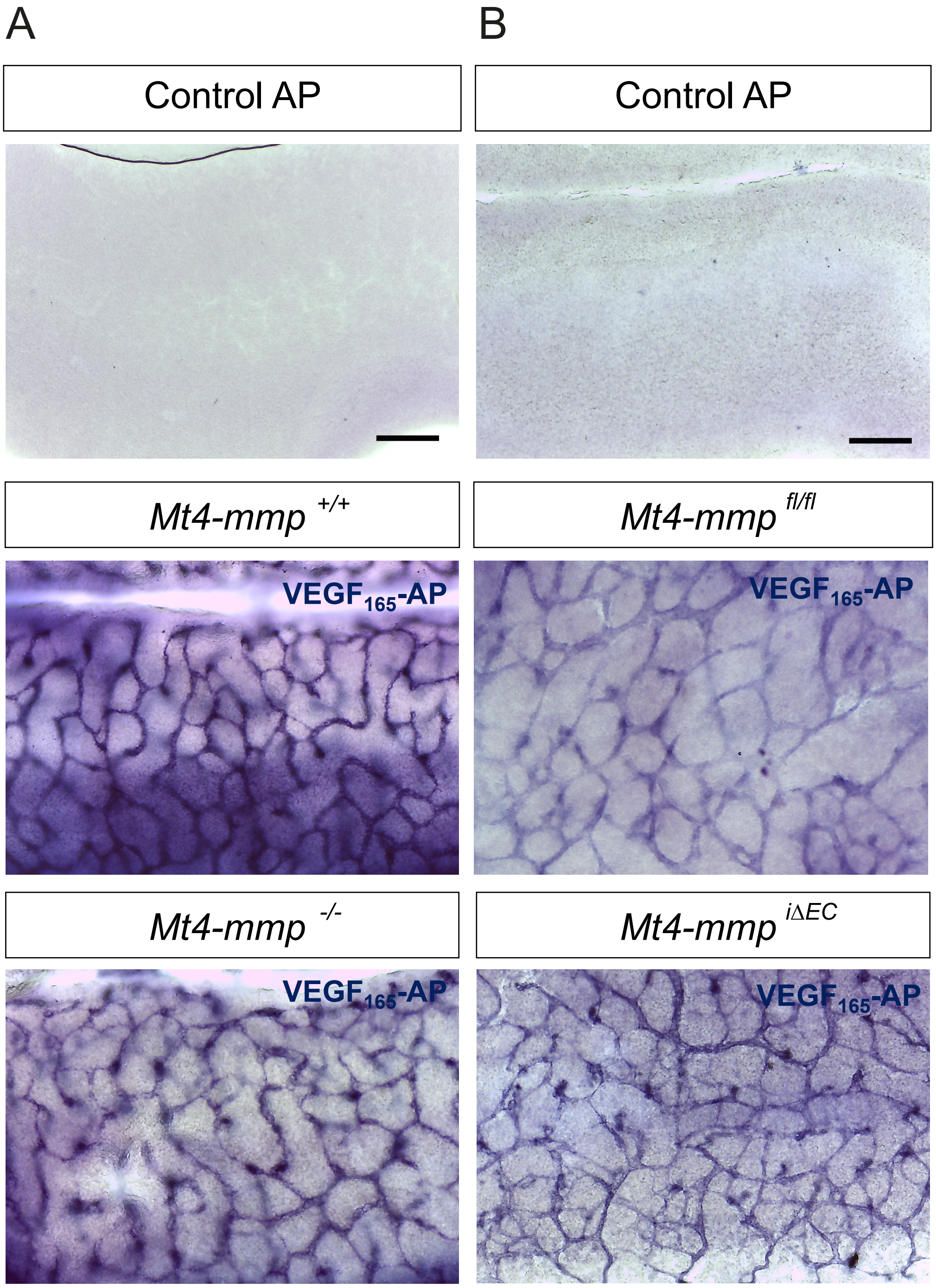
